# Simple Adjustment of Intra-nucleotide Base-phosphate Interaction in OL3 AMBER Force Field Improves RNA Simulations

**DOI:** 10.1101/2023.09.05.556403

**Authors:** Vojtěch Mlýnský, Petra Kührová, Petr Stadlbauer, Miroslav Krepl, Michal Otyepka, Pavel Banáš, Jiří Šponer

## Abstract

Molecular dynamics (MD) simulations represent an established tool to study RNA molecules. Outcome of MD studies depends, however, on the quality of the used force field (*ff*). Here we suggest a correction for the widely used AMBER OL3 *ff* by adding a simple adjustment of nonbonded parameters. The reparameterization of Lennard-Jones potential for the –H8…O5’– and –H6…O5’– atom pairs addresses an intra-nucleotide steric clash occurring in the type 0 base-phosphate interaction (0BPh). The non-bonded fix (NBfix) modification of 0BPh interactions (the NBfix_0BPh_ modification) was tuned via reweighting approach and, subsequently, tested using extensive set of standard and enhanced sampling simulations of both unstructured and folded RNA motifs. The modification corrects minor but visible intra-nucleotide clash for the *anti* nucleobase conformation. We observed that structural ensembles of small RNA benchmark motifs simulated with the NBfix_0BPh_ modification provide better agreement with experiments. No side-effects of the modification were observed in standard simulations of larger structured RNA motifs. We suggest that the combination of OL3 RNA *ff* and NBfix_0BPh_ modification is a viable option to improve RNA MD simulations.

## INTRODUCTION

Insights into RNA structural dynamics at atomistic description are essential for understanding of biomolecular motions and processes. Molecular dynamics (MD) simulation is an established theoretical tool that benefits from the ability to overcome experimental limits when finding links between RNA structure, dynamics, and function.^1-5^ Compact folded RNA molecules are typically well-described by modern empirical potentials (force fields, *ff*s) on a sub-μs timescale when starting simulations from established experimental structures. The general usage, applicability and reproducibility of MD simulations are continuously increasing. However, the MD simulation studies remain limited by the approximative nature of the *ff*s.

Number of RNA *ff*s are presently available to carry out MD simulations of RNA systems,^6-11^ but none of them is flawless.^5, 12-17^ Despite suboptimal performance for certain motifs, mainly the short RNA single strands,^14, 16, 18-20^ the AMBER OL3 *ff* from 2010^7^ still remains the state-of-the-art for RNA simulations and safest option to start with.^21^ It has been shown in the past decade that OL3 performance can be partially tweaked by optimizing simulation settings and by adding new *ff* terms that are orthogonal to the current *ff* terms. Namely, (i) an update of parameters for phosphate oxygens by Steinbrecher and coworkers^22^ was shown to improve the general simulation outcome,^23, 24^ (ii) combination of OL3 with the four-point OPC water model^25^ revealed benefits for structural description of short RNA single-strands (tetranucleotides, TNs) and tetraloops (TLs),^19, 24, 26^ (iii) controllable fine-tuning of specific pairwise H-bond interactions via external general H-bond fix potential (gHBfix) corrected structural ensembles of several RNA motifs,^13, 14, 27^ and (iv) specific adjustment of interactions formed by terminal nucleotides via tHBfix further improved the agreement with experiments for RNA TNs.^28^

In our recent work,^29^ we spotted another minor but visible imbalance of the AMBER *ff* connected with imprecise description of RNA intra-nucleotide interactions, when van der Waals (vdW) clash in G_S+1_(C8-H8)… G_S+1_(O5’) interaction resulted in the spurious flipping of the G_S+1_ sugar-phosphate backbone and population of nonnative structures of UUCG TL. The identified vdW clash in UUCG TL^29^ is related to the weak –CH…O– H-bond between –C8H8 and –C6H6 groups of purines and pyrimidines, respectively, and bridging O5’ phosphate oxygen from the same residue, which was categorized as base-phosphate interaction type 0 (0BPh, Figure 1).^30^ The attempt to adjust the 0BPh interaction in the structural context of UUCG TL via nonbonded fix (NBfix) stabilized the native G_S+1_ phosphate conformation^29^ and indicated that it could be beneficial for the stability of native states of both GAGA and UUCG TLs.^31^

**Figure 1:**
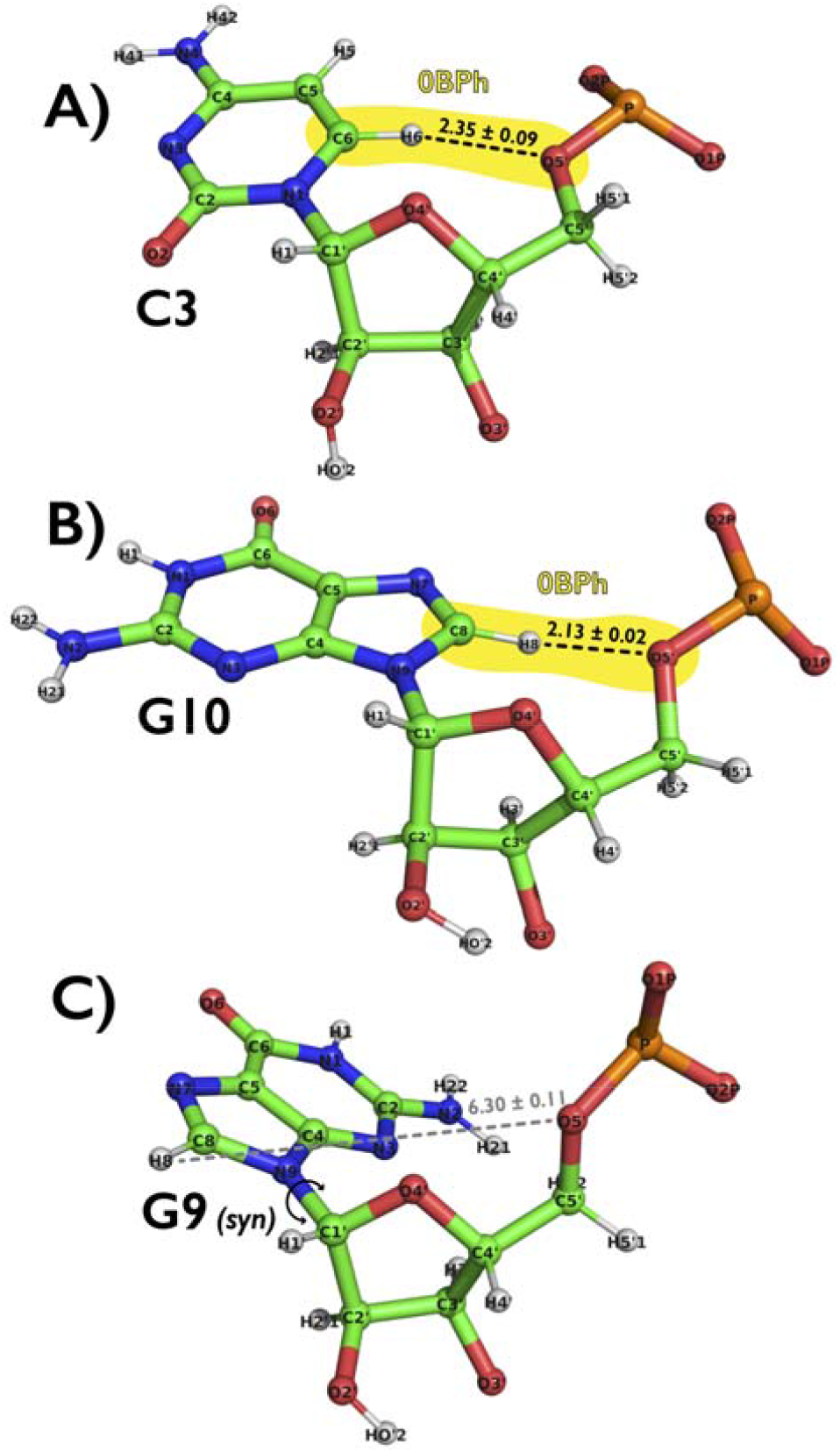
Definition and description of the weak –CH…O– H-bond that is classified as intra-nucleotide base-phosphate interaction type 0 (0BPh).^30^ Panels illustrate examples of 0BPh interactions with established –C6H6…O5’– and –C8H8…O5’– H-bonds for pyrimidines (C3, panel A) and purines (G10, panel B), respectively. The 0BPh is not established in *syn* orientation of nucleobase due to significantly higher –H6/H8…O5’– distances (G9, panel C). Snapshots of all three nucleotides were taken from the NMR structure of UUCG TL (PDB ID 2KOC)^38^ and measured distances between H6/H8 and O5’ atoms are shown as averages (with errors as standard deviations) over 20 deposited structures^38^.

In this work, we performed extensive set of standard and enhanced sampling MD simulations on diverse RNA systems and considered the adjustment of 0BPh interaction via NBfix (the NBfix_0BPh_ modification) as a general improvement of the AMBER OL3 RNA *ff* (and likely other *ff* variants using the original AMBER non-bonded parameters). We first used the reweighting approach to optimize settings for the NBfix_0BPh_ modification. Reweighting methods are one of the strongest tools of *ff* development,^26, 32^ and can re-evaluate the results of MD simulations under the assumption of a modified *ff* parametrization without repeating the entire simulation. Reweighting allowed us to identify most suitable modified parameters for both purines and pyrimidines, which were subsequently validated on benchmark systems during extensive set of MD simulations. The testing set contained single-stranded RNAs, duplexes, TLs and some examples of more structured RNA motifs with noncanonical interactions. We also investigated possible effects of the NBfix_0BPh_ modification (with similar setting as those for RNA nucleotides) on DNA duplex and guanine quadruplexes (GQs). We show that the NBfix_0BPh_modification improves structural behavior of small RNA motifs by increasing the agreement with experimental datasets. Stability of canonical A-RNA duplexes was also improved, whereas other, bigger, and more structured RNA motifs were not affected by the NBfix_0BPh_ modification on the affordable simulation timescale. In addition, this paper presents extensive set of new simulations with two versions of the external gHBfix potential and supports previous claims^14, 27^ that gHBfix is beneficial for proper structural description of RNA motifs.

## METHODS

### Details about AMBER RNA and DNA *ff*s and their adjustments

We used standard AMBER OL3 (known also as χOL3)^7, 33-35^ and AMBER OL15^36^ *ff*s for RNA and DNA simulations, respectively. OL3 RNA *ff* was further adjusted by the van der Waals (vdW) modification of phosphate oxygens developed by Steinbrecher et al.,^22^ where the affected dihedrals were adjusted as described elsewhere.^13, 23^ This RNA *ff* version is abbreviated as OL3_CP_ henceforth and AMBER library file can be found in Supporting Information of Ref. ^13^. In addition, we used gHBfix19 potential, which is the correction for the OL3_CP_ *ff* from 2019, where all –NH…N– base-base interactions are strengthened by 1.0 kcal/mol and all –OH…OR– and – OH…OR– sugar – phosphate interactions are weakened by 0.5 kcal/mol.^14^ We also tested the latest optimized gHBfix version from 2021 (gHBfix21), where all RNA H-bond donor…H-bond acceptor interactions are modified, i.e., base donor – base acceptor, base donor – sugar acceptor, base donor – phosphate acceptor, sugar donor – base acceptor, sugar donor – sugar acceptor, and sugar donor – phosphate acceptor are adjusted specifically (see Ref. ^27^ for full description). Simulations of RNA single strands were also performed with the tHBfix20 potential, where additional correction is added to interactions formed by terminal residues.^28^ For some specific tests, we also used standard OL3 (i.e., without phosphate modification) and older *ff*99bsc0^35^ RNA *ff*s.

On top of that, we modified the pairwise vdW parameters via breakage of the combination (mixing) rules via the nonbonded fix (NBfix) approach^37^ for atoms involved in 0BPh intra-nucleotide interactions for both RNA OL3_CP_ and DNA OL15 *ff*s. Namely, we reduced the minimum-energy distance of Lennard-Jones potential (i.e., *R*_*i,j*_ parameter) for the –H8…O5’– and –H6…O5’– pairs, i.e., between H5 – OR/OS and H4 – OR/OS atom types; by 0.25 Å to 2.8808 Å and 2.9308 Å for purine and pyrimidine nucleotides, respectively (modification labelled as NBfix_0BPh_). The OR atom type belongs to O5’ and O3’ bridging phosphate oxygens in the RNA OL3_CP_, whereas the OS atom type involves O5’, O3’ and also O4’ atoms from deoxyribose sugar in the DNA OL15 *ff*. Depths of the potential well (*ε*_*i,j*_ parameters) were kept at their default value of 0.0505 kcal/mol. In other words, we attempted to decrease repulsion between H8 and H6 atoms of all purine and pyrimidine bases and O5’ oxygens of phosphates. We note that O3’ and O4’ atoms have higher distances from H8 and H6 in comparison with those involving O5’ atoms and thus, modified Lennard-Jones potentials for H8/H6…O3’/O4’ atom pairs are supposed to have marginal effects.

### Starting structures and simulation setup

Initial coordinates of r(AAAA), r(CAAU), r(CCCC), r(GACC) and r(UUUU) tetranucleotides (TNs), r(UCAAUC) and r(UCUCGU) hexanucleotides (HNs) and r(gcGAGAgc) 8-mer tetraloop (GAGA TL) were prepared using Nucleic Acid Builder of AmberTools14^39^ as one strand of an A-form duplex. RNA single strands were solvated using a cubic box of the OPC water^25^ with a minimum distance between box walls and solute of 12 Å, yielding ∼2200 water molecules added (∼40×40×40 Å^3^ box size), ∼4100 water molecules added (∼50×50×50 Å^3^ box size), and ∼7600 water molecules added (∼60×60×60 Å^3^ box size) for TNs, HNs, and the GAGA TL, respectively. Enhanced sampling MD simulations were performed in ∼0.15 M of KCl salt (TNs and HNs) and ∼1.0 M KCl salt excess (GAGA TL) using Joung−Cheatham (JC) ionic parameters^40^ for the TIP4P-EW water.

Starting topologies and coordinates of two RNA duplexes (PDB ID 1QC0^41^ (only ten canonical base-pairs were considered) and 1RNA^42^), RNA Sarcin-Ricin loop (SRL, PDB ID 3DW4^43^; residues 2649-2671), RNA kink-turn 7 (Kt-7, PDB ID 1S72^44^; residues 76-83, 91-101), DNA duplex (PDB ID 1BNA^45^), and two guanine quadruplexes formed by the human telomeric sequence (GQ’s; parallel-stranded (PDB ID 1KF1^46^) and (3+1) hybrid (PDB ID 2GKU^47^)) were prepared from particular experimental structures by using the tLEaP module of AMBER 16 program package^48^ (see Supporting Information of Ref. ^14^ for details about structure preparation). Short r(CGCG)_2_ duplex was prepared using Nucleic Acid Builder of AmberTools14. Standard MD simulations were carried out in a cubic box of OPC^25^ and SPC/E^49^ water models (for RNA and DNA simulations, respectively) with a minimum distance between box walls and solute of 10 Å and with ∼0.15 M KCl salt using the JC ionic parameters. TIP3P water model^50^ was also used for some specific tests involving RNA duplexes (Table S1 in the Supporting Information).

All MD simulations were run at T = 298 K with the hydrogen mass repartitioning^51^ allowing an 4-fs integration time step (see Supporting Information of Ref. ^14^ for other details about minimization and equilibration protocols). Standard MD simulations were run in AMBER18,^52^ whereas both AMBER18 and GROMACS2018^53^ were used for enhanced sampling simulations. PARMED^54^ was used to convert AMBER topologies and coordinates into GROMACS inputs.

### Enhanced sampling simulations

We used two different enhanced sampling schemes, i.e., a standard replica exchange solute tempering (REST2) protocol^55^ and well-tempered metadynamics^56-58^ (MetaD) in combination with the REST2 method (ST-MetaD).^31, 59^ REST2 simulations were performed at 298 K (the reference replica) with 8 and 12 replicas for the TNs and HNs, respectively. The scaling factor (λ) values ranged from 1 to 0.601700871 and to 0.59984 for 8 and 12 replicas, respectively. Those values were chosen to maintain an exchange rate above 20%. The effective solute temperature ranged from 298 K to ∼500 K. REST2 simulations were performed with the AMBER GPU MD simulation engine (pmemd.cuda).^60^ Further details about REST2 settings can be found elsewhere.^14^ One test simulation of r(UCUCGU) HN was performed at 275 K temperature corresponding to the experimental conditions where the biggest number of NMR signals was obtained.^16^ The same λ values were applied for scaling, which resulted in effective solute temperature range from 275 K to ∼460 K. We note that the 275 K and 298 K REST2 simulations revealed comparable results considering limits of sampling.

ST-MetaD simulations of GAGA TL were performed with 12 replicas starting from unfolded single strands and were simulated in the effective temperature range of 298−497 K for 5 μs per replica. The average acceptance rate was ∼30%. The *ε*RMSD metric^61^ was used as a biased collective variable.^31, 62^ ST-MetaD simulations were carried out using a GPU-capable version of GROMACS2018^53^ in combination with PLUMED 2.5^63, 64^ (see Ref. ^31^ for further details about ST-MetaD settings). Besides newly performed MD simulations, we also used some trajectories from our previous works (see Table S1 in the Supporting Information for a full list of standard as well as enhanced sampling simulations).

The reweighting algorithm,^65^ which enables fast re-evaluation of the results from MD simulations, was used on the range of modified *R*_*i,j*_ values of the Lennard-Jones potential for the NBfix_0BPh_ modification in the attempt to find the optimal setup. Reweighting was performed on trajectories from both REST2 and ST-MetaD simulations using snapshots only from the reference replica (the lowest REST2 replica corresponding to 298 K; see Ref. ^31^ for further details about application of the reweighting approach).

### Conformational analysis

The most populated conformations from TNs and HNs structural ensembles were identified by clustering, which is based on an algorithm introduced by Rodriguez and Laio^66^ in combination with the εRMSD metric^61^ (see Ref. ^28^ for more details).

### Comparison between MD and NMR data

The conformational ensembles of TNs and HNs obtained from REST2 simulations were compared with previously published NMR experiments.^16, 20, 67-69^ We analyzed separately four NMR observables, i.e., (i) backbone ^3^J scalar couplings, (ii) sugar ^3^J scalar couplings, (iii) nuclear Overhauser effect intensities (NOEs), and (iv) the absence of specific peaks in NOE spectroscopy (uNOEs). In addition, we also considered ambiguous NOEs (ambNOEs) resulting from a sum of overlapping peaks^16, 20^ as fifth individual NMR component for HNs. ^3^J scalar couplings were calculated via Karplus relationships, NOEs and uNOEs were obtained as averages over the N samples, i.e.,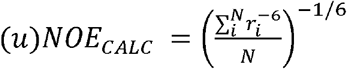, and ambNOEs were calculated by summing the contribution from either two or three nuclei pairs and again averaged over the N samples, i.e.,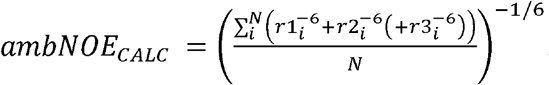 Combination of all those analyzed NMR observables (calculated as weighted arithmetic mean)provided the *total χ*^*2*^ value for each REST2 simulation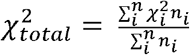 where 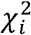and *n*_*i*_ are *individual χ*^*2*^ values and the number of observables, respectively, corresponding to the particular NMR observables. The lower *total χ*^*2*^ value, the better agreement between experiment and REST2 simulation (for detailed explanations, see, e.g., Refs. ^26, 28^).

ST-MetaD simulation of the GAGA TL provided populations of the native structure and other conformations, which can be used for estimation of the folding free energy balance (ΔGº_fold_).^27, 31^ Reference native structure of GAGA TL was taken from our previous work.^14^ The *ε*RMSD threshold separating the folded and un(mis)folded states was set at value of 0.7 (see Ref. ^31^ for details about ΔGº_fold_ estimations and convergence).

## RESULTS & DISCUSSION

### Reweighting reveals optimal settings for the NBfix_0BPh_ modification

Reweighting algorithm^65^ allows to efficiently re-evaluate results of MD simulations under the assumption of a modified *ff* parametrization without necessity to perform new simulations.^26, 31, 32^ Here, we reweighted NBfix_0BPh_ settings for a set of enhanced sampling simulations of RNA TNs and TLs. More specifically, we used results from OL3_CP_ REST2 simulations with gHBfix19 and tHBfix20 potentials of r(AAAA), r(CCCC), and r(CAAU) TNs^28^ and from folding OL3_CP_ ST-MetaD simulations with gHBfix19 of GAGA and UUCG TLs^31^. We monitored changes of total *χ*^*2*^ values (and its components) and ΔGº_fold_ energies for TNs and TLs, respectively. Each NBfix_0BPh_ setting was defined by scanning *R*_*i,j*_ values of the Lennard-Jones potential for the –H8…O5’– and –H6…O5’–atom pairs from 2.33 – 3.33 and 2.48 – 3.48 intervals for purine and pyrimidine nucleotides, respectively, using 0.025 Å steps.

Results from TN simulations indicate that 0.25 Å decrease of *R*_*i,j*_ parameters for both purine and pyrimidine nucleotides is optimal (Figures S1-S5 in Supporting Information). Reweighted data from r(CCCC) and r(CAAU) simulations with the sole gHBfix19 suggest even larger decrease (than the finally selected 0.25 Å) of the *R*_*i,j*_ value for pyrimidines, i.e., –H6…O5’–atom pairs, but that is contradicted by data from simulations with combined gHBfix19+tHBfix20 potentials (those giving much better agreement with experiments;^28^ see Figures S2, S4 and S5 in Supporting Information). Data from reweighting of ST-MetaD TL simulations support results from TNs. Reweighting of the GAGA TL simulation shows that adjustment of *R*_*i,j*_ values for the –H8…O5’– atom pairs of purine nucleotides is the dominant contributor for shifting ΔGº_fold_ energy (Figure 2A). This is not surprising since six out of eight nucleotides from r(gcGAGAgc) TL are purines. Data even suggested that larger decrease would be vital, but that was not confirmed when reweighting *R*_*i,j*_ values for both purines and pyrimidines simultaneously (Figure 2A).

**Figure 2:**
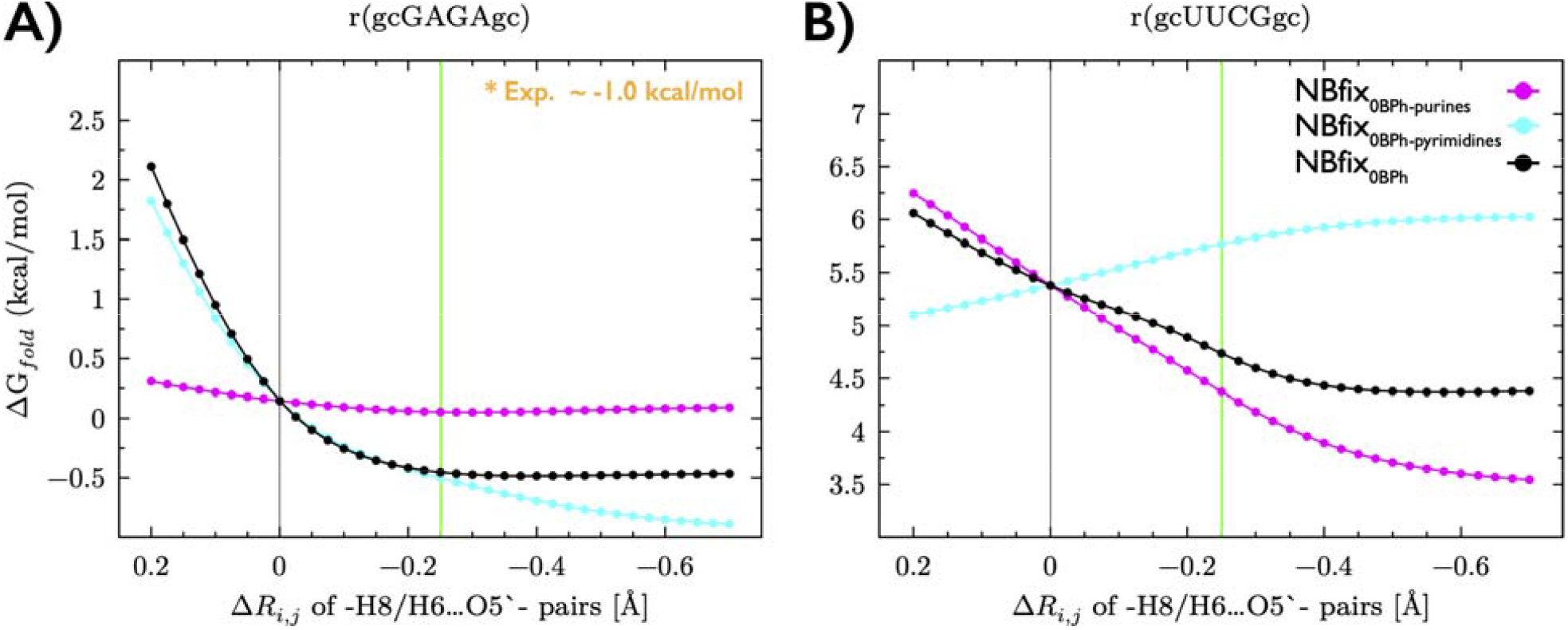
Effects of different setting of NBfix_0BPh_ modifications on stability of the native state of RNA TLs. Folding free energies (ΔGº_fold_) were estimated by reweighting of GAGA (panel A) and UUCG (panel B) trajectories from OL3_CP_ ST-MetaD simulations with gHBfix19 potential. Plots show dependence of ΔGº_fold_ energies on modified vdW parameters for purines and pyrimidines, i.e., changes of *R*_*i,j*_ values for –H8…O5’– (cyan line) and –H6…O5’– (magenta line) atom pairs from the original (standard AMBER *ff*) values. Black line presents the outcome, where *R*_*i,j*_ values for both purines and pyrimidines were modified simultaneously by the same value (see Methods for details). Gray vertical line indicates position of the original (unmodified) values. The NBfix_0BPh_ modification (0.25 Å decrease of pairwise terms; green vertical line) stabilizes native states of both TLs by providing lower ΔGº_fold_ energies, which are in better agreement with experiments (reported ΔGº_fold_ energies are between -0.7 and -1.3 kcal/mol for both TLs^70-72^).

Modifications of *R*_*i,j*_ parameters for both purines and pyrimidines affect significantly the stability of UUCG TL native state, but in opposite direction (Figure 2B). More specifically, decrease of the *R*_*i,j*_ value for only pyrimidine residues indicates stabilization; ΔGº_fold_ is decreasing from ∼5.4 kcal/mol (default *R*_*i,j*_ value) up to ∼3.5 kcal/mol (final and lowest tested *R*_*i,j*_), Figure 2B. In contrary, decreased *R*_*i,j*_ values for sole purines gives destabilization (ΔGº_fold_ is increasing from ∼5.4 kcal/mol up to ∼6.0 kcal/mol, Figure 2B). Such a counterintuitive effect of the decreased *R*_*i,j*_ values for the –H8…O5’– atom pairs of purines, i.e., small destabilization of the UUCG native state, is caused by stabilization of misfolded states, where the key G residue in the loop (G_L4_) is sampling noncanonical *anti* orientation of χ dihedral (*syn* states are required for the native conformation and those are not affected by the NBfix_0BPh_ modification; see Figure 1 and Ref. ^31^ for further discussion). Most importantly, simultaneous modification of *R*_*i,j*_ values for both purines and pyrimidines, i.e., the NBfix_0BPh_ modification, revealed substantial stabilization of the UUCG native state (i.e., modified pyrimidines provide dominant effect for the UUCG TL, Figure 2B).

In summary, reweighted data from both TN and TL motifs showed that decreased *R*_*i,j*_ parameters for both –H8…O5’– and –H6…O5’–atom pairs provided better agreement with experiments. Effective usage of reweighting requires broad exploration of the conformational space, which we believe was achieved in trajectories from REST2 and ST-MetaD simulations of TNs and TLs, respectively. The accuracy of reweighted results decreases with increasing deviation of the parameter from original values (see, e.g., continuous decrease of ΔGº_fold_ energy with decreasing *R*_*i,j*_ parameter for pyrimidines on Figure 2B). Those large parameter changes often result in problems (e.g., clashes) in subsequent simulations with modified parameters as they drive system towards states that were not sampled in original trajectories used for reweighting. Hence, we aimed for reasonably small change of *R*_*i,j*_ parameters that would be enough to relax the steric clash of –H8…O5’– and –H6…O5’–atom pairs observed in the original OL3_CP_ *ff*. We found that 0.25 Å decrease of the particular *R*_*i,j*_ parameter for both purines and pyrimidines appears to be the optimal choice.

### NBfix_0BPh_ modification of the AMBER OL3_CP_ RNA *ff* improves structural description of single-stranded RNA motifs

We used the optimized setting for the NBfix_0BPh_ modification (0.25 Å decrease) and explicitly tested its effect on structural description of five RNA TNs and two HNs, i.e., benchmark set of motifs with available NMR data, in a large set of enhanced sampling simulations. We again opted for the REST2 protocol, which was shown to provide ensembles with sufficient convergence for those single-stranded motifs.^14, 28^ We used OL3_CP_ *ff* combined with gHBfix19, gHBfix19+tHBfix20 and gHBfix21+tHBfix20 potentials, which were shown to improve structural behavior of the short RNA single strands motifs (see Methods and Table S1 in Supporting Information). The data is summarized in Tables 1 and 2 for TNs and HNs, respectively.

**Table 1:** REST2 simulations of RNA TNs and their comparison with experiments. ^a^

| Motif | gHBfix version | NBfix<sub>0BPh</sub> | $\chi^2$<br>(<sup>3</sup>J-backbone, <sup>3</sup>J-sugar, NOE,<br>uNOE (# of violations) / **Total**) <sup>b</sup> | Clustering<br>(A-form / Loop / Bulge-out<br>/ Intercalated; in %) | # of<br>clusters |
| --- | --- | --- | --- | --- | --- |
| r(GACC) <sup>c</sup> | gHBfix19 | No | 0.39, 0.50, 1.13, 0.34 (3) / **0.40** | ~74/~7/~5/~2 | 5 |
| r(GACC) <sup>c</sup> | gHBfix19_tHBfix20 | No | 0.34, 0.37, 1.11, 0.15 (3) / **0.23** | ~81/~5/~4/~0 | 4 |
| r(GACC) | gHBfix21_tHBfix20 | No | 0.28, 0.39, 2.62, 0.02 (1) / **0.20** | ~86/~2/~4/~0 | 3 |
| r(GACC) | gHBfix19 | Yes | 0.29, 0.33, 1.23, 0.00 (0) / **0.10** | ~91/~1/~3/~0 | 4 |
| r(GACC) | gHBfix19_tHBfix20 | Yes | 0.29, 0.34, 1.34, 0.00 (1) / **0.11** | ~90/~2/~4/~0 | 4 |
| r(GACC) | gHBfix21_tHBfix20 | Yes | 0.28, 0.39, 2.70, 0.00 (0) / **0.19** | ~93/~2/~2/~0 | 3 |
| r(CAAU) <sup>c</sup> | gHBfix19 | No | 0.73, 1.14, 1.81, 6.07 (28) / **5.37** | ~33/~0/~3/~51 | 5 |
| r(CAAU) <sup>c</sup> | gHBfix19 + tHBfix20 | No | 0.49, 0.80, 1.40, 2.22 (29) / **2.05** | ~76/~0/~6/~3 | 8 |
| r(CAAU) | gHBfix21 + tHBfix20 | No | 0.47, 1.55, 1.63, 3.78 (36) / **3.41** | ~73/~0/~9/~1 | 5 |
| r(CAAU) | gHBfix19 | Yes | 0.55, 0.92, 1.28, 4.88 (21) / **4.30** | ~49/~0/~0/~44 | 3 |
| r(CAAU) | gHBfix19 + tHBfix20 | Yes | 0.33, 0.56, 0.89, 0.32 (12) / **0.37** | ~89/~0/~0/~5 | 3 |
| r(CAAU) | gHBfix21 + tHBfix20 | Yes | 0.30, 0.51, 1.02, 0.65 (12) / **0.66** | ~85/~0/~6/~0 | 5 |
| r(AAAA) <sup>c</sup> | gHBfix19 | No | 0.66, 1.23, 1.08, 1.14 (12) / **1.11** | ~30/~0/~6/~10 | 17 |
| r(AAAA) <sup>c</sup> | gHBfix19 + tHBfix20 | No | 0.68, 1.48, 1.28, 1.06 (11) / **1.08** | ~32/~0/~5/~1 | 18 |
| r(AAAA) | gHBfix21 + tHBfix20 | No | 0.61, 1.84, 1.07, 0.82 (6) / **0.87** | ~38/~0/~8/~0 | 15 |
| r(AAAA) | gHBfix19 | Yes | 0.44, 0.83, 1.10, 0.38 (5) / **0.48** | ~66/~0/~3/~12 | 7 |
| r(AAAA) | gHBfix19 + tHBfix20 | Yes | 0.42, 0.78, 1.14, 0.02 (2) / **0.20** | ~72/~0/~3/~2 | 8 |
| r(AAAA) | gHBfix21 + tHBfix20 | Yes | 0.35, 0.66, 1.21, 0.12 (2) / **0.28** | ~70/~0/~8/~0 | 5 |
| r(CCCC) <sup>c</sup> | gHBfix19 | No | 0.44, 0.68, 1.54, 3.22 (11) / **2.83** | ~55/~0/~0/~40 | 2 |
| r(CCCC) <sup>c</sup> | gHBfix19 + tHBfix20 | No | 0.25, 0.38, 1.22, 0.50 (6) / **0.55** | ~85/~0/~2/~8 | 5 |
| r(CCCC) | gHBfix21 + tHBfix20 | No | 0.22, 0.35, 3.38, 0.10 (3) / **0.41** | ~86/~0/~5/~3 | 4 |
| r(CCCC) | gHBfix19 | Yes | 0.28, 0.38, 1.28, 1.87 (5) / **1.68** | ~76/~0/~0/~23 | 2 |
| r(CCCC) | gHBfix19 + tHBfix20 | Yes | 0.20, 0.30, 1.41, 0.12 (3) / **0.25** | ~93/~0/~0/~4 | 2 |
| r(CCCC) | gHBfix21 + tHBfix20 | Yes | 0.18, 0.27, 3.92, 0.02 (1) / **0.39** | ~96/~0/~0/~0 | 1 |
| r(UUUU) <sup>c</sup> | gHBfix19 | No | 0.53, 0.87, 3.53, 1.65 (19) / **1.63** | ~5/~0/~0/~3 | 10 |
| r(UUUU) <sup>c</sup> | gHBfix19 + tHBfix20 | No | 0.52, 0.97, 3.46, 1.86 (17) / **1.81** | ~14/~0/~0/~0 | 10 |
| r(UUUU) | gHBfix21 + tHBfix20 | No | 0.47, 1.69, 3.58, 1.47 (11) / **1.49** | ~8/~25/~0/~0 | 9 |
| r(UUUU) | gHBfix19 | Yes | 0.48, 0.54, 4.50, 1.72 (19) / **1.70** | ~36/~0/~2/~2 | 9 |
| r(UUUU) | gHBfix19 + tHBfix20 | Yes | 0.49, 0.60, 4.55, 2.04 (18) / **1.99** | ~29/~0/~5/~0 | 9 |
| r(UUUU) | gHBfix21 + tHBfix20 | Yes | 0.46, 1.06, 4.44, 1.71 (12) / **1.71** | ~24/~27/~1/~0 | 11 |

<sup>a</sup> Simulations used combination of OL3<sub>CP</sub> RNA *ff* with either gHBfix19 or gHBfix21 potentials and sometimes also with tHBfix20 (see Methods). REST2 simulations used 8 replicas and were run for 10 μs (the last 7 μs were used for data analysis). Experimental data are taken from Refs. <sup>67-69</sup>.
<sup>b</sup> $\chi^2$ values were obtained by comparing calculated and experimental backbone <sup>3</sup>J scalar couplings, sugar <sup>3</sup>J scalar couplings, nuclear Overhauser effect intensities (NOEs), and the absence of specific peaks in NOE spectroscopy (uNOEs, number of violations is also reported). See Methods for details.
<sup>c</sup> Data taken from previous papers. <sup>14, 28</sup>

**Table 2:**
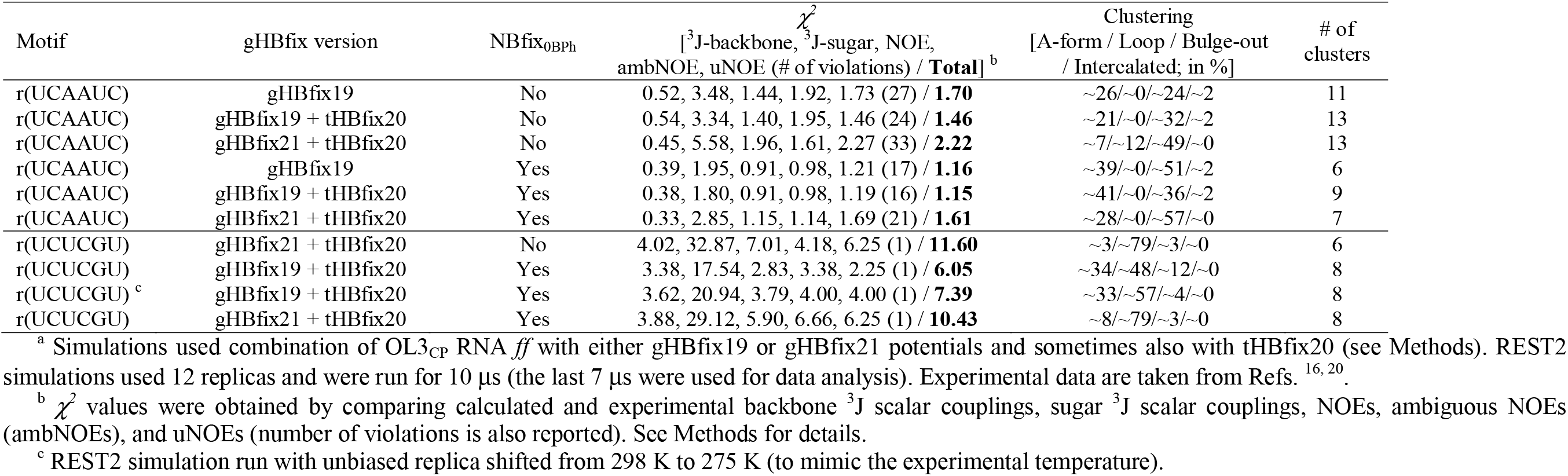
REST2 simulations of RNA HNs and their comparison with experiments. ^a^.

The NBfix_0BPh_ modification further fine-tunes especially r(CAAU) and r(AAAA) ensembles, which now agree excellently with NMR data (total χ^2^ value below 1; Table 1). The NBfix_0BPh_ modification indirectly stabilizes *anti* state of A residues and thus resolves previously reported excessive sampling of *syn* states.^28, 73^ We also observed that the NBfix_0BPh_ increased population of canonical A-form like states in structural ensembles of all simulated TNs (Table 1), which is expected to be the main conformer based on available experimental data.^20, 67-69^

Application of the NBfix_0BPh_ did not provide better agreement with the experiment for r(UUUU) TN. We did observe increase (∼20%) in population of the canonical A-form like structures (as for other TNs), but it was associated with slight increase of total χ^2^ values, i.e., agreement between simulations and experiment was modestly worsened for all tested MD setups (Table 1). Experimental data shows out that r(UUUU) is more dynamical than others TNs and should sample unstacked/disordered states.^67, 74^ Hence, further adjustment of AMBER RNA *ff*s is probably required potentially including revision of stacking interactions,^73^ in order to improve behavior of poly-U sequences in MD simulations.

Recently, modification of AMBER potential (named STAfix) by major weakening of intra-molecular stacking was employed to study spontaneous binding of RNA single strands to RNA recognition motif proteins.^17^ It efficiently eliminated majority of previously identified spurious r(UUUU) 1-3_2-4 stacked conformers^28^ and supported sampling of unstructured states in structural ensembles, which resulted in excellent agreement with the NMR data.^17^ STAfix, however, is not suitable for simulations of folded RNAs; it is a special-purpose modification which has been prepared for binding of unstructured single-stranded RNAs to proteins. We also note that we have significantly less NMR signals for r(UUUU) than for other TNs, which complicates predictions from the experiment.

The NBfix_0BPh_ modification improves structural behavior of two r(UCAAUC) and r(UCUCGU) HN motifs. It has been shown previously by both experiments and simulations that r(UCAAUC) should sample predominantly structures around the canonical A-form state.^20, 75^ On the other hand, presence of G residue in r(UCUCGU) HN significantly affects the conformational space and this HN motif is much more dynamical than shorter RNA TNs.^16^ r(UCAAUC) REST2 simulations performed here show that the NBfix_0BPh_ modification slightly enhanced population of canonical A-form like states for r(UCAAUC) HN and results are in good agreement with the experiment^20^ (total χ^2^ values close to 1; Table 2). r(UCUCGU) REST2 simulations revealed disagreement with experiment with significantly higher total χ^2^ values than obtained for other tested single-stranded motifs. As the NBfix_0BPh_ modification did not appear to significantly affect r(UCUCGU) structural ensembles, we performed simulations with only four different setups for this motif (Table 2). The gHBfix21 potential provided slightly worse agreement with the experiment^16^ than gHBfix19 as it increased population of loop-like structures. We also performed one REST2 simulation at lower temperature (where the biggest number of NMR signals was obtained, see Methods), but it did not improve the agreement (Table 2). We made some initial attempt to identify the source of this huge discrepancy between experiment and theory (see the next section).

In summary, the NBfix_0BPh_ modification affects positively structural ensembles of single-stranded RNA motifs. Most importantly, it resolves previously reported excessive sampling of *syn* states for A residues^28, 73^ by indirectly stabilizing *anti* states. r(UUUU) TN and especially r(UCUCGU) HN motifs remain challenging for the AMBER OL3_CP_ *ff* even with the additional refinements. Despite some uncertainty in experimental datasets for those motifs, it is evident that larger reparameterization than simple adjustment of pairwise terms for one interaction would be required to improve structural behavior of those motifs. Taking all the available data together clearly demonstrates that we remain far from having flawless *ff* for RNA molecules.

### Measured NMR signals for r(UCUCGU) HN are from multiple structures

r(UCUCGU) REST2 simulations revealed striking disagreement between MD predictions and the experimental data.^16^ We observed that calculated χ^2^ values are almost order of magnitude higher than those from TN simulations and the other r(UCAAUC) HN (Tables 1 and 2). Inspection of individual χ^2^ components (see Methods) revealed that largest deviations between experimentally measured and predicted results from simulations come from sugar ^3^J scalar couplings.

Experimental data suggest that each residue is sampling mixture of C3’-endo and C2’-endo sugar puckers.^16^ Indeed, MD is able to sample transitions between these most common RNA pucker conformations but overall populations are shifted from those measured experimentally for all residues (with G5 residue showing the largest deviation; see Table S2 in Supporting Information). Analysis of most populated conformers during REST2 simulation showed that besides the canonical A-form-like states, MD favors formation of a loop-like conformer with formation of G5C2 base pair (Figure S6 in Supporting Information). The experiment shows much weaker NMR A-form signals than for TN and r(UCAAUC) HN motifs which indicates presence of another state.^16^ However, formation of conformers with G5C2 base pair is not supported.^16^

In summary, r(UCUCGU) NMR data are composed of different conformers but those sampled during REST2 simulations (besides the canonical A-form-like states) are apparently different from those observed in the experiment. We speculate that a different preference for the C2’-endo sugar pucker in the AMBER OL3_CP_ *ff* simulation could also (to some extent) affect the structural ensemble of the r(UCUCGU) HN. This is out of scope of this work and will be addressed in future.

### The native state of GAGA TL is stabilized by the NBfix_0BPh_ modification

We performed ST-MetaD simulation of r(gcGAGAgc) TL to explicitly test the effect of the NBfix_0BPh_ modification on folding of this small 8-mer RNA TL motif. We obtained ΔGº_fold_ energy of -0.5 ± 0.6 kcal/mol (corresponding to 68.3 ± 20.0 % population of the native state). The result is in an excellent agreement with data from the reweighting approach (see Figure 2 and Ref. ^31^). Despite the fact that the “true” uncertainty in convergence of ST-MetaD simulations is, probably, larger that the statistical error (and the results should be ideally derived by series of independent simulations^31, 76, 77^), the agreement between predicted data from reweighting and the actual simulation result is encouraging. The GAGA native state is stabilized by ∼0.6 kcal/mol in comparison with the control simulation (ΔGº_fold_ = 0.1 ± 0.2 kcal/mol)^31^ indicating that the NBfix_0BPh_ modification could increase the population of the GAGA native state by ∼24% at 298 K.

### MD simulations of common NA motifs with the NBfix_0BPh_ modification did not reveal any side-effects

Despite clear benefits of the NBfix_0BPh_ modification on structural description of small RNA motifs, one should always test its effects on bigger and structured RNA and DNA motifs before any claims are made about its general applicability. Hence, we performed standard MD simulations of systems commonly used for NA *ff* validation, i.e., RNA and DNA duplexes, RNA SRL and Kt-7 motifs, and two DNA GQs. We compared results with the control simulation set (i.e., the same *ff* setup just without the NBfix_0BPh_; see Methods). We observed that the NBfix_0BPh_ modification did not introduce any side effects in structural description of these motifs (full details are in Supporting Information).

### The NBfix_0BPh_ modification disfavors spurious ladder-like structures

We showed in previous section that the NBfix_0BPh_ supports canonical A-form backbone conformation in short RNA single strands. In addition, simulations of RNA duplexes revealed that similarly to OL3 correction,^78^ the NBfix_0BPh_ modestly reduces an inclination of the A-form duplex (see Supporting Information). Therefore we also tested possible effect of the NBfix_0BPh_ on potential formation of spurious structure form termed as a ladder-like RNA structure,^79^ which is associated with population of high-*anti* χ angle in RNA duplexes. Elimination of the ladder-like structure was the key improvement introduced by the OL3 parametrization.^7^ Thus, we tested the NBfix_0BPh_ on longer 1QC0 duplex^41^ (i.e., r(GCACCGUUGG)_2_ decamer, see Methods) and short r(CGCG)_2_ duplex known for a quick (∼10 ns) transition from the canonical A-form to the spurious ladder-like structure in obsolete AMBER *ff* versions lacking the OL3 correction.^78^ We compared behavior of two *ff*s, i.e., the outdated *ff*99bsc0^35^ and the currently recommended OL3 *ff*, in combination with two different water models (TIP3P^50^ and OPC^25^), and explicitly probed possible effects of adding the NBfix_0BPh_ modification. As a result, we compared behavior of eight different simulation setups (Table S3 in Supporting Information) on both duplexes. The initial set of r(CGCG)_2_ simulations revealed high tendency to fraying, i.e., base-pair opening of terminal base pairs, which significantly affected outcomes of the simulations (Table S3 in Supporting Information). Therefore, we used the gHBfix21 potential^27^ in the next set of r(CGCG)_2_ simulations to stabilize base-base interactions and suppress the extensive base pair fraying. We observed that a spontaneous transition into the ladder structure occurred only in the *ff*99bsc0+TIP3P setup for both short and long duplexes (Table S3 in Supporting Information). Thus, besides the known reparameterization of χ dihedrals of nucleobases within the OL3 parameterization^7^, there might be other factors affecting propensity of transitions of canonical duplexes to ladders, i.e., (i) the four-point OPC water model, and (ii) the NBfix_0BPh_ modification (effective removal of the clash in the sugar-phosphate backbone^29^). Subsequently, we started simulations from the ladder conformation of the r(CGCG)_2_ duplex and observed that only setups with OL3 *ff* were able to undergo successful transitions back to the canonical A-form structures on the 1 μs timescale; the ladders were corrected in all four cases on timescales ∼100 – 250 ns. The NBfix_0BPh_ alone did not eliminate the ladder, at least on 1 μs timescale (Table S3 in Supporting Information).

In summary, the NBfix_0BPh_ modification is a vital addition to the standard AMBER OL3 RNA *ff*. It can further stabilize canonical A-form duplexes and prevent transitions into artificial ladders. Results from duplex simulations revealed that *anti*/high-*anti* imbalance of nucleotides (corrected by crucial OL3 parameterization^7^) was not the only driving factor for spurious transitions into ladder-like structures. The clash in the backbone remained^29^ even after applying OL3 correction and this clash can be efficiently removed by the NBfix_0BPh_ modification. The OL3 *ff* can be safely combined with the NBfix_0BPh_ and this combination further reduces the likelihood of sampling various spurious structures.

## CONCLUDING REMARKS

We presented the adjustment of pairwise Lennard-Jones parameters of the OL3 AMBER RNA *ff* for atoms involved in the 0BPh intra-nucleotide molecular interactions, i.e., the NBfix_0BPh_ modification. This was motivated by the recently identified spurious steric clash in the sugar-phosphate backbone associated with the 0BPh interaction.^29^ We optimized NBfix_0BPh_ settings by the reweighting method, which is an efficient approach to assess effects of *ff* changes without the necessity to perform actual extensive simulations. Reweighting data indicate that the 0.25 Å decrease of pairwise *R*_*i,j*_ parameters for both –H8…O5’– and –H6…O5’–atom pairs is the optimal adjustment.

The NBfix_0BPh_ modification was further verified by extensive set of new enhanced sampling simulations for contemporary recommended set of benchmark RNA systems including tetranucleotides, hexanucleotides and tetraloops. The simulations with the NBfix_0BPh_ modification provided better agreement with available experimental data. We suggest that the NBfix_0BPh_ modification should be used in all future simulations of these small RNA oligonucleotides since it directly corrects a real imbalance of the AMBER *ff* non-bonded terms. Elimination of its consequences by other *ff* terms like dihedral potential reparameterizations or gHBfix is thus not desirable. We also did not find any undesired side-effects of the NBfix_0BPh_ modification in standard simulations of larger and more complicated RNA motifs with noncanonical interactions. The data for canonical duplexes indicate that NBfix_0BPh_ may help to prevent A-RNA distortions into the ladder-like structure, though the OL3 still remains essential in preventing this artifact.

Hence, the NBfix_0BPh_ modification could be considered as a general addition to the OL3 RNA *ff* (and likely also for other *ff* variants using the original AMBER non-bonded parameters). The NBfix_0BPh_ provides healthier sampling of nucleotides in *anti* conformation. Considering results shown here, we suggest the utilization of the NBfix_0BPh_ modification on top of the AMBER OL3 *ff*,^7^ modified parameters for phosphates by Steinbrecher et al.,^22^ OPC water model,^25^ and the gHBfix21 potential,^27^ for MD simulations of RNA especially for small motifs such as short single strands and tetraloops.

In summary, NBfix_0BPh_ modification corrects a moderate but evident imbalance in the repulsion term of the AMBER *ff* non-bonded terms for the 0BPh interaction, which affects the accuracy mainly in case of the simulations of small RNA oligonucleotides. As the NBfix_0BPh_ modification reparametrizes only one off-diagonal Lennard-Jones parameter between different atom types, its application is selective, robust and without any side-effects, so it can be used in combination with any *ff* from the AMBER family.

## Supporting information

Supporting Information to the Article

## ASSOCIATED CONTENT

### Supporting Information

The Supporting Information is available free of charge via the Internet at http://pubs.acs.org/ and containing details about standard MD simulations of various RNA and DNA motifs, supporting tables and figures to the article (PDF). Coordinates of the starting systems are also attached (zipped PDBs).

## AUTHOR INFORMATION

## ACKNOWLEDGMENT

This work was supported by the Czech Science Foundation to VM, PS, MK and JS (grant number 23-05639S). This research also received the support of EXA4MIND, a European Union’s Horizon Europe Research and Innovation programme under grant agreement N° 101092944. Views and opinions expressed are however those of the author(s) only and do not necessarily reflect those of the European Union or the European Commission. Neither the European Union nor the granting authority can be held responsible for them.

