## Supporting Information to the Article for "Simple Adjustment of Intra-nucleotide Base-phosphate Interaction in OL3 AMBER Force Field Improves RNA Simulations"

### SUPPORTING RESULTS

**Standard MD simulations of RNA and DNA motifs with the NBfix_0BPh_ modification show no negative side-effects.** We performed a set of unbiased MD simulations to investigate effects of the NBfix_0BPh_ modification on a range of RNA systems and common DNA motifs (Table S1). We used the OL3_CP_ and OL15 force fields (*ff*s) for RNA and DNA simulations, respectively (see Methods in the main text). The OL3_CP_ RNA *ff* was corrected by the gHBfix19 potential^1^ (all –NH…N– base-base interactions are strengthened and all –OH…OR– and –OH…OR– sugar – phosphate interactions are weakened; see Methods in main text for details). We also used the gHBfix19 for DNA duplex simulations, where only –NH…N– base-base interactions were strengthened in the same way as for RNA simulations (sugar – phosphate interactions were not affected due to missing –OH groups).

Initially, we simulated two RNA duplexes, i.e., r(GCACCGUUGG)_2_ decamer (excised from the PDB ID 1QC0^2^ structure) and r(UUAUAUAUAUAUAA)_2_ tetradecamer (PDB ID 1RNA^3^). We performed two independent 5 μs-long standard MD simulations of each above-mentioned duplex with the NBfix_0BPh_ modification and compared them with the control simulations, i.e., those with the same settings just without the NBfix_0BP_ as control runs. Duplex simulations with the NBfix_0BPh_ modification showed stable behavior, which resulted in similar averaged RMSD values (calculated from the starting structures) as those obtained from control simulations (1.1 ± 0.2 Å and 1.6 ± 0.4 Å for the decamer and tetradecamer, respectively). We observed decreased base-pair opening frequency of both GC pairs and AU base pairs at the end of helices in simulations with NBfix_0BPh_ modification. Terminal base-pair breathing (fraying) was reduced by ~2% and ~11% for dodecamer (GC pairs) and tetradecamer (AU pairs), respectively (Table S4). We note that observed fraying results could be within the sampling limits of the simulations and that accurate (unambiguous) experimental data for the quantification of frayed structures are still not available for RNA sequences. However, the decreased probability of terminal base-pair fraying is expected to have positive effects on the behavior of RNA systems in MD simulations (see Refs. ^1, 4^ for further discussion about base-pair fraying). The analysis of helical base-pair parameters, i.e., those measuring orientation and displacement of Watson–Crick base pairs, showed that the NBfix_0BPh_ did not affect simulated A-RNA duplexes. Averaged values of helical parameters from control and NBfix_0BPh_-modified simulations are mostly within statistical errors (showing fluctuations, Table S4). The NBfix_0BPh_ modification has visible effect on calculated inclination and base pair roll of both RNA duplexes, where we identified modest decreases of averaged values by ~1.6° and 0.6° for the inclination and roll parameters, respectively; Table S4). Note that the helical structure parameter inclination and base pair step parameter roll are mathematically interconnected.^5^ In summary, the NBfix_0BPh_ has minimal effect on the A-RNA structure and the terminal base-pairs of A-RNA duplexes appear to be slightly more stable.

Next, we performed two independent 5 μs-long simulations of Drew-Dickerson DNA dodecamer (d(CGCGAATTCGCG)_2_, PDB ID 1BNA^6^) in both settings, i.e., with and without the NBfix_0BPh_ modification. DNA duplex also revealed stable behavior upon the addition of NBfix_0BPh_ with similar averaged RMSD as the control set of simulations (1.5 ± 0.3 Å). Helical base-pair parameters obtained from simulations with the NBfix_0BPh_ modification and control simulations are entirely comparable (Table S5). The NBfix_0BPh_ modification did result in noticeable change of the propeller parameter (averaged values are increased by ~0.9° in comparison with control simulations), but those differences are much smaller than statistical errors showing fluctuations (Table S5).

We then performed two independent 5 μs-long simulations of the kink-turn Kt-7 motif (see Methods in main text and Figure S7A) with the NBfix_0BPh_ modification and compared them with prolonged simulations from our previous work^1^, i.e., simulations without the NBfix_0BPh_ used as control sets (Table S1). We observed the H-bonds characterizing A-minor I interaction,^7^ i.e., G6(2’-OH)…A16(N1), C11(2’-OH)…A16(O2’), and G6(N2H)…A16(N3), were lost in one simulation with the NBfix_0BPh_ modification (~3 µs) and somewhat earlier during both control simulations (before ~1 us, Figures S8 and S9). In all three simulations, this loss was immediately followed by transition into A-minor 0 interaction, which is characterized by formation of G6(2’-OH)…A16(N3) and G6(O2’)…A16(2’-OH) H-bonds (Figures S8 and S9). Transitions from A-minor I to A-minor 0 are well-known and occurring regularly in Kt-7 simulations, when this kink-turn is simulated out of its structural context.^8^ We note the reversible attempts to transition from A-minor I into A-minor 0 were also observed during second simulation with the NBfix_0BPh_ modification (Figure S9). In conclusion, the dynamics and stability of the Kt-7 motif was qualitatively same in NBfix_0BPh_-modified and control simulations.

We then tested the NBfix_0BPh_ modification on the Sarcin-Ricin loop RNA motif (see Methods in main text and Figure S7B). The overall SRL fold remained stable and we observed comparable dynamics between simulations with the NBfix_0BPh_ modification and control runs (one prolonged simulation from our previous work^1^ and one entirely new simulation, Table S1). We identified that the stability of one BPh contact, i.e., the G16(N1)…U8(O2P) H-bond, is weakened (presumably water-bridged) for most of the time in both NBfix_0BPh_-modified and control simulation sets (Figures S10 and S11). Importantly, the structural stability of both the whole GpU platform, i.e., residues G7, U8 and A17, and the neighboring trans Hoogsteen/Sugar Edge A9G16 base pair were unaffected (Figures S10 and S11). Thus, SRL simulations with and without the NBfix_0BPh_ modification provided basically identical outcome within the limits of sampling.

Finally, we investigated possible influence of the NBfix_0BPh_ modification on structural behavior of the parallel-stranded (1KF1)^9^ and (3+1) hybrid (2GKU)^10^ DNA GQ’s during 2 μs-long standard MD simulations. We observed very similar (basically identical within the limits of sampling) structural behavior of both tested GQ’s in simulations with and without the NBfix_0BPh_. Both cation binding sites inside the GQ channel were fully occupied in all the simulations. The distributions of G-quartet planarities, stacking distances and helical twists showed insignificant differences between the tested *ff* variants (Figures S12 and S13). Population of the state with bifurcated hydrogen bonds in G-quartets^1^ differed by a few per cents between the *ff*s, but without a uniform trend, so we assume it was caused by the stochastic nature of MD simulations (Table S6). Altogether, the results indicate no effect of the NBfix_0BPh_ modification on GQ simulation behavior.

In summary, no adverse structural side-effects of the NBfix_0BPh_ modification were noticed in our testing set of standard simulations of common folded RNA and DNA motifs.

### SUPPORTING TABLES

**Table S1:** Overview of all performed enhanced sampling and standard MD simulations.^a^

| RNA Systems | *ff* | WAT | gHBfix | NBfix_0BPh_ | Method | Reps. ^b^ | Length [μs] |
| --- | --- | --- | --- | --- | --- | --- | --- |
| r(GACC) ^c^ | OL3_CP_ | OPC | gHBfix19 | No | REST2 | 8 | 10 |
| r(GACC) ^d^ | OL3_CP_ | OPC | gHBfix19, tHBfix20 | No | REST2 | 8 | 10 |
| r(GACC) | OL3_CP_ | OPC | gHBfix21, tHBfix20 | No | REST2 | 8 | 10 |
| r(GACC) | OL3_CP_ | OPC | gHBfix19 | Yes | REST2 | 8 | 10 |
| r(GACC) | OL3_CP_ | OPC | gHBfix19, tHBfix20 | Yes | REST2 | 8 | 10 |
| r(GACC) | OL3_CP_ | OPC | gHBfix21, tHBfix20 | Yes | REST2 | 8 | 10 |
| r(CAAU) ^c^ | OL3_CP_ | OPC | gHBfix19 | No | REST2 | 8 | 10 |
| r(CAAU) ^d^ | OL3_CP_ | OPC | gHBfix19, tHBfix20 | No | REST2 | 8 | 10 |
| r(CAAU) | OL3_CP_ | OPC | gHBfix21, tHBfix20 | No | REST2 | 8 | 10 |
| r(CAAU) | OL3_CP_ | OPC | gHBfix19 | Yes | REST2 | 8 | 10 |
| r(CAAU) | OL3_CP_ | OPC | gHBfix19, tHBfix20 | Yes | REST2 | 8 | 10 |
| r(CAAU) | OL3_CP_ | OPC | gHBfix21, tHBfix20 | Yes | REST2 | 8 | 10 |
| r(AAAA) ^c^ | OL3_CP_ | OPC | gHBfix19 | No | REST2 | 8 | 10 |
| r(AAAA) ^d^ | OL3_CP_ | OPC | gHBfix19, tHBfix20 | No | REST2 | 8 | 10 |
| r(AAAA) | OL3_CP_ | OPC | gHBfix21, tHBfix20 | No | REST2 | 8 | 10 |
| r(AAAA) | OL3_CP_ | OPC | gHBfix19 | Yes | REST2 | 8 | 10 |
| r(AAAA) | OL3_CP_ | OPC | gHBfix19, tHBfix20 | Yes | REST2 | 8 | 10 |
| r(AAAA) | OL3_CP_ | OPC | gHBfix21, tHBfix20 | Yes | REST2 | 8 | 10 |
| r(CCCC) ^c^ | OL3_CP_ | OPC | gHBfix19 | No | REST2 | 8 | 10 |
| r(CCCC) ^d^ | OL3_CP_ | OPC | gHBfix19, tHBfix20 | No | REST2 | 8 | 10 |
| r(CCCC) | OL3_CP_ | OPC | gHBfix21, tHBfix20 | No | REST2 | 8 | 10 |
| r(CCCC) | OL3_CP_ | OPC | gHBfix19 | Yes | REST2 | 8 | 10 |
| r(CCCC) | OL3_CP_ | OPC | gHBfix19, tHBfix20 | Yes | REST2 | 8 | 10 |
| r(CCCC) | OL3_CP_ | OPC | gHBfix21, tHBfix20 | Yes | REST2 | 8 | 10 |
| r(UUUU) ^c^ | OL3_CP_ | OPC | gHBfix19 | No | REST2 | 8 | 10 |
| r(UUUU) ^d^ | OL3_CP_ | OPC | gHBfix19, tHBfix20 | No | REST2 | 8 | 10 |
| r(UUUU) | OL3_CP_ | OPC | gHBfix21, tHBfix20 | No | REST2 | 8 | 10 |
| r(UUUU) | OL3_CP_ | OPC | gHBfix19 | Yes | REST2 | 8 | 10 |
| r(UUUU) | OL3_CP_ | OPC | gHBfix19, tHBfix20 | Yes | REST2 | 8 | 10 |
| r(UUUU) | OL3_CP_ | OPC | gHBfix21, tHBfix20 | Yes | REST2 | 8 | 10 |
| r(UCAAUC) | OL3_CP_ | OPC | gHBfix19 | No | REST2 | 12 | 10 |
| r(UCAAUC) | OL3_CP_ | OPC | gHBfix19, tHBfix20 | No | REST2 | 12 | 10 |
| r(UCAAUC) | OL3_CP_ | OPC | gHBfix21, tHBfix20 | No | REST2 | 12 | 10 |
| r(UCAAUC) | OL3_CP_ | OPC | gHBfix19 | Yes | REST2 | 12 | 10 |
| r(UCAAUC) | OL3_CP_ | OPC | gHBfix19, tHBfix20 | Yes | REST2 | 12 | 10 |
| r(UCAAUC) | OL3_CP_ | OPC | gHBfix21, tHBfix20 | Yes | REST2 | 12 | 10 |
| r(UCUCGU) | OL3_CP_ | OPC | gHBfix21, tHBfix20 | No | REST2 | 12 | 10 |
| r(UCUCGU) | OL3_CP_ | OPC | gHBfix19, tHBfix20 | Yes | REST2 | 12 | 10 |
| r(UCUCGU) ^e^ | OL3_CP_ | OPC | gHBfix19, tHBfix20 | Yes | REST2 | 12 | 10 |
| r(UCUCGU) | OL3_CP_ | OPC | gHBfix21, tHBfix20 | Yes | REST2 | 12 | 10 |
| gcGAGAgc ^f^ | OL3_CP_ | OPC | gHBfix19 | No | ST-MetaD | 12 | 5 |
| gcGAGAgc | OL3_CP_ | OPC | gHBfix19 | Yes | ST-MetaD | 12 | 5 |
| gcUUCGgc ^f^ | OL3_CP_ | OPC | gHBfix19 | No | ST-MetaD | 12 | 5 |
| 1QC0 | OL3_CP_ | OPC | gHBfix19 | No | sMD | 1 | 5 |
| 1QC0 | OL3_CP_ | OPC | gHBfix19 | Yes | sMD | 2 | 5 |
| 1RNA | OL3_CP_ | OPC | gHBfix19 | No | sMD | 1 | 5 |
| 1RNA | OL3_CP_ | OPC | gHBfix19 | Yes | sMD | 2 | 5 |
| Kt-7 ^g^ | OL3_CP_ | OPC | gHBfix19 | No | sMD | 2 | 5 |
| Kt-7 | OL3_CP_ | OPC | gHBfix19 | Yes | sMD | 2 | 5 |
| SRL ^g^ | OL3_CP_ | OPC | gHBfix19 | No | sMD | 2 | 5 |
| SRL | OL3_CP_ | OPC | gHBfix19 | Yes | sMD | 2 | 5 |
| r(CGCG)_2_ | *ff*99bsc0 | TIP3P | - | No | sMD | 1 | 1 |
| r(CGCG)_2_ | *ff*99bsc0 | TIP3P | - | Yes | sMD | 1 | 1 |
| r(CGCG)_2_ | *ff*99bsc0 | OPC | - | No | sMD | 1 | 1 |
| r(CGCG)_2_ | *ff*99bsc0 | OPC | - | Yes | sMD | 1 | 1 |
| r(CGCG)_2_ | OL3 | TIP3P | - | No | sMD | 1 | 1 |
| r(CGCG)_2_ | OL3 | TIP3P | - | Yes | sMD | 1 | 1 |
| r(CGCG)_2_ | OL3 | OPC | - | No | sMD | 1 | 1 |
| r(CGCG)_2_ | OL3 | OPC | - | Yes | sMD | 1 | 1 |
| r(CGCG)_2_ | *ff*99bsc0 | TIP3P | gHBfix21 | No | sMD | 1 | 1 |
| r(CGCG)_2_ | *ff*99bsc0 | TIP3P | gHBfix21 | Yes | sMD | 1 | 1 |
| r(CGCG)_2_ | *ff*99bsc0 | OPC | gHBfix21 | No | sMD | 1 | 1 |
| r(CGCG)_2_ | *ff*99bsc0 | OPC | gHBfix21 | Yes | sMD | 1 | 1 |
| r(CGCG)_2_ | OL3 | TIP3P | gHBfix21 | No | sMD | 1 | 1 |
| r(CGCG)_2_ | OL3 | TIP3P | gHBfix21 | Yes | sMD | 1 | 1 |
| r(CGCG)_2_ | OL3 | OPC | gHBfix21 | No | sMD | 1 | 1 |
| r(CGCG)_2_ | OL3 | OPC | gHBfix21 | Yes | sMD | 1 | 1 |
| 1QC0 | *ff*99bsc0 | TIP3P | - | No | sMD | 1 | 1 |
| 1QC0 | *ff*99bsc0 | TIP3P | - | Yes | sMD | 1 | 1 |
| 1QC0 | *ff*99bsc0 | OPC | - | No | sMD | 1 | 1 |
| 1QC0 | *ff*99bsc0 | OPC | - | Yes | sMD | 1 | 1 |
| 1QC0 | OL3 | TIP3P | - | No | sMD | 1 | 1 |
| 1QC0 | OL3 | TIP3P | - | Yes | sMD | 1 | 1 |
| 1QC0 | OL3 | OPC | - | No | sMD | 1 | 1 |
| 1QC0 | OL3 | OPC | - | Yes | sMD | 1 | 1 |
| r(CGCG)_2_ | *ff*99bsc0 | TIP3P | gHBfix21 | Yes | sMD | 1 | 1 |
| r(CGCG)_2_ | *ff*99bsc0 | OPC | gHBfix21 | No | sMD | 1 | 1 |
| r(CGCG)_2_ | *ff*99bsc0 | OPC | gHBfix21 | Yes | sMD | 1 | 1 |
| r(CGCG)_2_ | OL3 | TIP3P | gHBfix21 | No | sMD | 1 | 1 |
| r(CGCG)_2_ | OL3 | TIP3P | gHBfix21 | Yes | sMD | 1 | 1 |
| r(CGCG)_2_ | OL3 | OPC | gHBfix21 | No | sMD | 1 | 1 |
| r(CGCG)_2_ | OL3 | OPC | gHBfix21 | Yes | sMD | 1 | 1 |
| DNA Systems | *ff* | WAT | gHBfix | NBfix_0BPh_ | Method | Reps.^b^ | Length [μs] |
| 1BNA | OL15 | SPC/E | gHBfix19 ^h^ | No | sMD | 2 | 5 |
| 1BNA | OL15 | SPC/E | gHBfix19 ^h^ | Yes | sMD | 2 | 5 |
| 1KF1 | OL15 | SPC/E | - | No | sMD | 1 | 2 |
| 1KF1 | OL15 | SPC/E | - | Yes | sMD | 1 | 2 |
| 2GKU | OL15 | SPC/E | - | No | sMD | 1 | 2 |
| 2GKU | OL15 | SPC/E | - | Yes | sMD | 1 | 2 |

^a^ See Methods in main text for details about system and *ff* labelling, description of HBfix potentials and setting of the NBfix_0BPh_ modification.

^b^ Number of replicas in enhanced sampling simulations (REST2 / ST-MetaD) or number of independent simulations for standard MD (sMD) simulations.

^c^ Data taken from Ref. ^1^.

^d^ Data taken from Ref. ^11^.

^e^ REST2 simulation run with unbiased replica shifted from 298 K to 275 K (to mimic the experimental temperature).

^f^ Data taken from Ref. ^12^.

^g^ One (SRL motif) or two (Kt-7 motif) 1 μs-long MD simulations from Ref. ^1^ were prolonged to 5 μs. We performed one entirely new independent simulation of the SRL motif.

^h^ The –NH…N– base-base interactions were stabilized by 1.0 kcal/mol (i.e., identically to gHBfix19 for RNA; destabilization of sugar-phosphate H-bonding is not applicable for DNA).

**Table S2:** Population of the C2’-endo pucker of r(UCUCGU) HN residues from simulations and experiment.^a^

| r(UCUCGU) | Fraction of C2’-endo-like sugar pucker | |
| --- | --- | --- |
|  | REST2 | Exp. ^b^ |
| U1 | 0.21 | 0.25 |
| C2 | 0.13 | 0.25 |
| U3 | 0.62 | 0.37 |
| C4 | 0.59 | 0.28 |
| G5 | 0.12 | 0.55 |
| U6 | 0.50 | 0.43 |

^a^ Data taken from REST2 simulation with OL3_CP_, gHBfix19 and tHBfix20 potentials and with the NBfix_0BPh_ modification (see Methods in main text and Table S1).

^b^ Experimental data measured at 275 K.^13^

**Table S3:** Occurrence of spurious ladder-like structures in simulations of RNA duplexes.^a^

| System | *ff* version | Correction | Water model | Fraying, top pair [%] ^b^ | Fraying, bottom pair [%] ^b^ | Ladder [ns] |
| --- | --- | --- | --- | --- | --- | --- |
| CGCG | *ff*99bsc0 | - | TIP3P | ~24 | ~35 | ~7 |
|  | *ff*99bsc0 | - | OPC | ~70 | ~53 | ~12 |
|  | *ff*99bsc0 | NBfix_0BPh_ | TIP3P | ~95 | ~93 | ~13 – ~49 ^c^ |
|  | *ff*99bsc0 | NBfix_0BPh_ | OPC | ~43 | ~75 | ~45 – ~85, ~335 |
|  | OL3 | - | TIP3P | ~57 | ~25 | - |
|  | OL3 | - | OPC | ~94 | ~26 | ~764 |
|  | OL3 | NBfix_0BPh_ | TIP3P | ~47 | ~10 | - |
|  | OL3 | NBfix_0BPh_ | OPC | ~27 | ~16 | - |
| CGCG | *ff*99bsc0 | gHBfix21 | TIP3P | ~0 | ~1 | ~504 |
|  | *ff*99bsc0 | gHBfix21 | OPC | ~0 | ~0 | - |
|  | *ff*99bsc0 | gHBfix21+NBfix_0BPh_ | TIP3P | ~33 | ~7 | - |
|  | *ff*99bsc0 | gHBfix21+NBfix_0BPh_ | OPC | ~0 | ~0 | - |
|  | OL3 | gHBfix21 | TIP3P | ~0 | ~0 | - |
|  | OL3 | gHBfix21 | OPC | ~0 | ~3 | - |
|  | OL3 | gHBfix21+NBfix_0BPh_ | TIP3P | ~0 | ~0 | - |
|  | OL3 | gHBfix21+NBfix_0BPh_ | OPC | ~1 | ~0 | - |
| 1QC0 | *ff*99bsc0 | - | TIP3P | ~0 | ~83 | ~150 |
|  | *ff*99bsc0 | - | OPC | ~3 | ~10 | - |
|  | *ff*99bsc0 | NBfix_0BPh_ | TIP3P | ~0 | ~2 | - |
|  | *ff*99bsc0 | NBfix_0BPh_ | OPC | ~1 | ~9 | - |
|  | OL3 | - | TIP3P | ~0 | ~0 | - |
|  | OL3 | - | OPC | ~65 | ~1 | - |
|  | OL3 | NBfix_0BPh_ | TIP3P | ~0 | ~0 | - |
|  | OL3 | NBfix_0BPh_ | OPC | ~0 | ~0 | - |
| CGCG | *ff*99bsc0 ^d^ | gHBfix21 | OPC | ~0 | ~0 | - |
|  | *ff*99bsc0 ^d^ | gHBfix21+NBfix_0BPh_ | TIP3P | ~1 | ~1 | - |
|  | *ff*99bsc0 ^d^ | gHBfix21+NBfix_0BPh_ | OPC | ~0 | ~0 | - |
|  | OL3 ^d^ | gHBfix21 | TIP3P | ~9 | ~2 | ~98 ^e^ |
|  | OL3 ^d^ | gHBfix21 | OPC | ~14 | ~0 | ~167 ^e^ |
|  | OL3 ^d^ | gHBfix21+NBfix_0BPh_ | TIP3P | ~7 | ~1 | ~237 ^e^ |
|  | OL3 ^d^ | gHBfix21+NBfix_0BPh_ | OPC | ~0 | ~1 | ~189 ^e^ |

^a^ All simulations were run for 1 μs (see Methods in main text and Table S1).

^b^ Top and bottom pairs are denoted to residues C1G8 and G4C5 for r(CGCG)_2_ duplex and G1C20 and G10C11 for 1QC0 duplex, respectively.

^c^ We observed unfolding, i.e., strand separation, of the duplex structure at ~557 ns.

^d^ Simulations were initiated form the ladder-like structure.

^e^ Time denotes complete transition back to the canonical A-form duplex, i.e., reparation of the ladder-like structure.

**Table S4:** Averaged values of helical parameters (and standard deviations represented by error bars) and base pair fraying of 1QC0 and 1RNA duplexes.^a^

|  | Inclination [°] | Slide [°] | Twist [°] | Rise [Å] | Tip [°] | Tilt [°] | Shift [Å] | Buckle [°] | Opening [°] | Shear [Å] | Stagger [Å] | Stretch [Å] | Propeller [°] | Roll [°] | Minor groove [Å] | Major groove [Å] | Fraying (top, bottom) ^b^ [%] | |
| --- | --- | --- | --- | --- | --- | --- | --- | --- | --- | --- | --- | --- | --- | --- | --- | --- | --- | --- |
| 1QC0 (Exp.) | 15.2 | -1.7 | 30.8 | 2.7 | 2.1 | -1.2 | -0.1 | 1.2 | 1.4 | 0.1 | 0.1 | -0.3 | -12.5 | 8.1 | 13.1 | 18.1 | - | - |
| gHBfix19 | 14.3  ± 0.7 | -1.8 ± 0.1 | 31.3 ± 0.4 | 2.7 ± 0.0 | -0.6 ± 0.3 | 0.3 ± 0.1 | 0.0 ± 0.0 | 1.5 ± 0.9 | 0.1 ± 0.2 | 0.0 ± 0.1 | -0.1 ± 0.0 | 0.0 ± 0.0 | -12.3 ± 0.7 | 7.7 ± 0.4 | 13.4 ± 0.0 | 19.2 ± 0.0 | 4.2 | 0.1 |
| gHBfix19 + NBfix_0BPh_ | 12.9  ± 0.5 | -1.6 ± 0.1 | 31.2 ± 1.2 | 2.8  ± 0.1 | -0.5 ± 0.1 | 0.3 ± 0.1 | 0.0 ± 0.0 | 1.3 ± 0.2 | -0.2 ± 0.1 | 0.0 ± 0.0 | -0.3 ± 0.0 | 0.0 ± 0.0 | -13.7 ± 0.5 | 7.3 ± 0.3 | 13.4 ± 0.5 | 18.7 ± 0.7 | 0.1 | 0.0 |
|  | 12.9  ± 0.5 | -1.6 ± 0.6 | 31.3 ± 1.2 | 2.8 ± 0.1 | -0.5 ± 0.1 | 0.3 ± 0.0 | 0.0 ± 0.0 | 1.3 ± 0.2 | -0.3 ± 0.1 | 0.0 ± 0.0 | -0.2 ± 0.0 | 0.0 ± 0.0 | -13.8 ± 0.6 | 7.4 ± 0.3 | 13.4 ± 0.5 | 18.7 ± 0.7 | 0.1 | 0.1 |
| 1RNA(Exp.) | 18.8 | -1.3 | 30.6 | 2.6 | -0.4 | 0.1 | 0.1 | 1.3 | -0.4 | -0.1 | 0.0 | -0.2 | -18.8 | 10.0 | 13.3 | 18.3 | - | - |
| gHBfix19 | 18.1  ± 0.5 | -1.5 ± 0.0 | 30.2 ± 0.1 | 2.5 ± 0.0 | 0.0 ± 0.1 | 0.0 ± 0.1 | 0.0 ± 0.0 | 0.0 ± 0.2 | 0.8 ± 0.2 | 0.0 ± 0.0 | 0.1 ± 0.0 | 0.0 ± 0.0 | -17.5 ± 0.2 | 10.5 ± 0.3 | 13.3 ± 0.0 | 19.2 ± 0.0 | 22.4 | 7.1 |
| gHBfix19 + NBfix_0BPh_ | 16.4  ± 0.6 | -1.3 ± 0.1 | 31.6 ± 1.2 | 2.6 ± 0.1 | 0.0 ± 0.1 | 0.0 ± 0.0 | 0.0 ± 0.0 | 0.0 ± 0.1 | 0.9 ± 0.1 | 0.0 ± 0.0 | 0.1 ± 0.0 | 0.0 ± 0.0 | -17.3 ± 0.7 | 9.7 ± 0.4 | 13.3 ± 0.5 | 18.7 ± 0.7 | 2.5 | 3.1 |
|  | 16.4  ± 0.6 | -1.3 ± 0.1 | 31.5 ± 1.2 | 2.6 ± 0.1 | 0.0 ± 0.1 | 0.0 ± 0.0 | 0.0 ± 0.0 | 0.0 ± 0.2 | 0.9 ± 0.1 | 0.0 ± 0.0 | 0.1 ± 0.0 | 0.0 ± 0.0 | -17.3 ± 0.7 | 9.8 ± 0.4 | 13.3 ± 0.5 | 18.7 ± 0.7 | 4.6 | 3.7 |

^a^ All simulation were run in OL3_CP_ *ff*, OPC water model and 0.15 M KCl (see Methods in main text and Table S1). The first two base pairs from each termini were excluded to prevent end effects and standard deviations were calculated using block average over 1000 snapshots (saved every 10 ps).

^b^ Top and bottom pairs are denoted to residues G1C20 and G10C11 for 1QC0 duplex and U1A28 and A14U15 for 1RNA duplex, respectively.

**Table S5:** Averaged values of helical parameters (and standard deviations represented by error bars) and base pair fraying of 1BNA duplex.^a^

|  | Inclination [°] | Slide [°] | Twist [°] | Rise [Å] | Tip [°] | Tilt [°] | Shift [Å] | Buckle [°] | Opening [°] | Shear [Å] | Stagger [Å] | Stretch [Å] | Propeller [°] | Roll [°] | Minor groove [Å] | Major groove [Å] | Fraying (top, bottom) ^b^ [%] | |
| --- | --- | --- | --- | --- | --- | --- | --- | --- | --- | --- | --- | --- | --- | --- | --- | --- | --- | --- |
| 1BNA (Exp.) | 0.4 | 0.1 | 35.6 | 3.3 | 0.0 | -0.2 | -0.1 | -0.7 | 0.1 | 0.1 | 0.1 | -0.2 | -13.6 | -0.3 | 12.3 | 18.9 | - | - |
| gHBfix19 | 2.5  ± 5.1 | 0.0 ± 0.4 | 35.7 ± 2.9 | 3.3 ± 0.1 | -0.1 ± 3.5 | 0.0 ± 2.1 | 0.0 ± 0.4 | 0.0 ± 4.6 | -0.4 ± 0.7 | 0.0 ± 0.1 | 0.0 ± 0.1 | -0.0 ± 0.0 | -10.1 ± 7.3 | 1.3 ± 2.9 | 12.2 ± 0.2 | 19.2 ± 0.3 | 1.7 | 60.6 |
|  | 2.6  ± 5.2 | 0.0 ± 0.4 | 35.6 ± 3.1 | 3.3 ± 0.1 | 0.0 ± 3.1 | 0.0 ± 1.9 | 0.0 ± 0.4 | 0.0 ± 4.6 | -0.5 ± 0.7 | 0.0 ± 0.1 | 0.1 ± 0.1 | 0.0 ± 0.0 | -10.8 ± 6.1 | 1.3 ± 2.9 | 12.2 ± 0.2 | 19.2 ± 0.4 | 0.1 | 0.8 |
| gHBfix19 + NBfix_0BPh_ | 1.8  ± 5.3 | 0.0 ± 0.4 | 35.8 ± 3.5 | 3.3 ± 0.1 | 0.1 ± 3.7 | -0.1 ± 2.3 | 0.0 ± 0.4 | -0.1 ± 4.8 | -0.5 ± 0.8 | 0.0 ± 0.1 | 0.1 ± 0.1 | 0.0 ± 0.0 | -9.5 ± 7.6 | 0.8 ± 2.9 | 12.2 ± 0.2 | 19.1 ± 0.4 | 69.2 | 1.3 |
|  | 1.8  ± 5.3 | 0.0 ± 0.4 | 35.8 ± 3.5 | 3.3 ± 0.1 | -0.1 ± 3.9 | 0.0 ± 2.3 | 0.0 ± 0.4 | 0.1 ± 4.8 | -0.5 ± 0.8 | 0.0 ± 0.1 | 0.1 ± 0.1 | 0.0 ± 0.0 | -9.6 ± 7.4 | 0.8 ± 2.9 | 12.2 ± 0.2 | 19.1 ± 0.4 | 0.1 | 62.0 |

^a^ All simulation were run in OL15 *ff*, SPC/E water model and 0.15 M KCl (see Methods in main text and Table S1). The first two base pairs from each termini were excluded to prevent end effects and standard deviations were calculated using block average over 1000 snapshots (saved every 10 ps).

^b^ Top and bottom pairs are denoted to residues C1G24 and G12C13, respectively.

**Table S6:** Population fraction of normal, bifurcated and other states of the middle quartet observed in the simulations of 1KF1 and 2GKU GQs.

| Simulation | Normal state | Bifurcated state | Other state |
| --- | --- | --- | --- |
| 1KF1 | 0 | 0.948 | 0.052 |
| 1KF1, NBfix_0BPh_ | 0 | 0.918 | 0.082 |
| 2GKU | 0 | 0.948 | 0.052 |
| 2GKU, NBfix_0BPh_ | 0 | 0.961 | 0.039 |

^a^ All simulation were run in OL15 *ff*, SPC/E water model and 0.15 M KCl (see Methods in main text and Table S1).

### SUPPORTING FIGURES

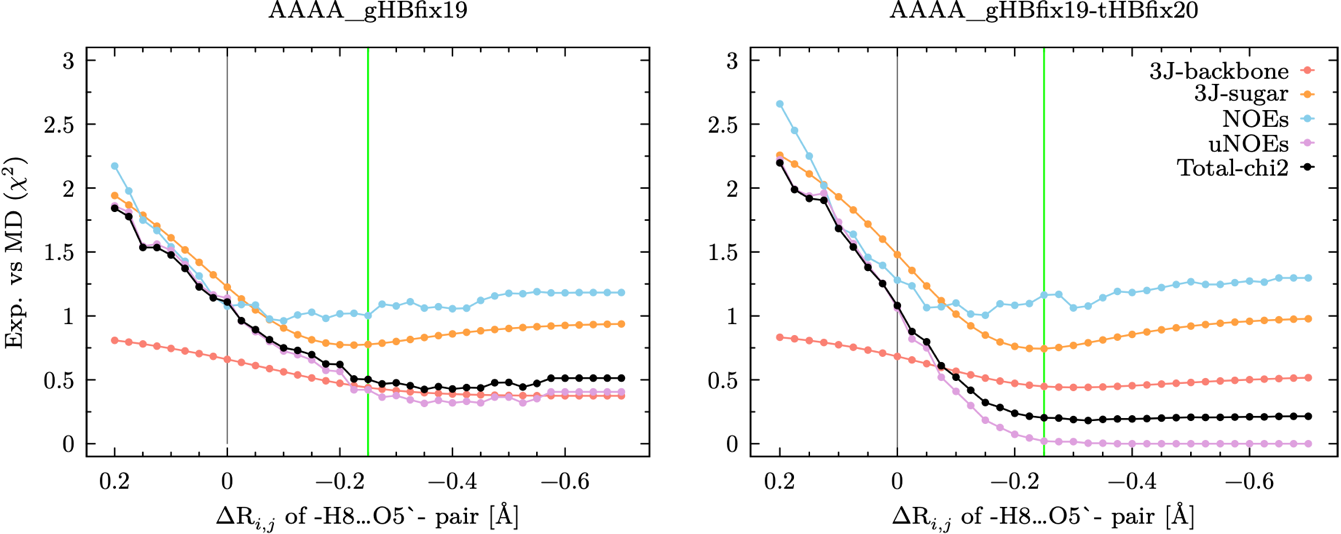

**Figures S1:** Effects of different settings of the NBfix_0BPh_ modification on agreement between simulations and NMR data as probed by reweighting of two r(AAAA) TN trajectories from REST2 simulations (only trajectories from reference, i.e., unbiased, replicas were considered for the analysis). Evolution of NMR observables, i.e., ^3^J scalar couplings, NOEs, uNOEs, and total χ^2^ values (combination of all observables; see Methods in main text) from OL3_CP_ REST2 simulations with gHBfix19 (panel A) and gHBfix19+tHBfix20 (panel B). Plots show dependence of NMR observables on changes of the Lennard-Jones *R_i,j_* parameter for purines, i.e., the distance between –H8…O5’– pairs, from the original (standard AMBER *ff*) values. Grey and green vertical lines mark positions of original and modified (i.e., chosen for further testing) parameter values, respectively.

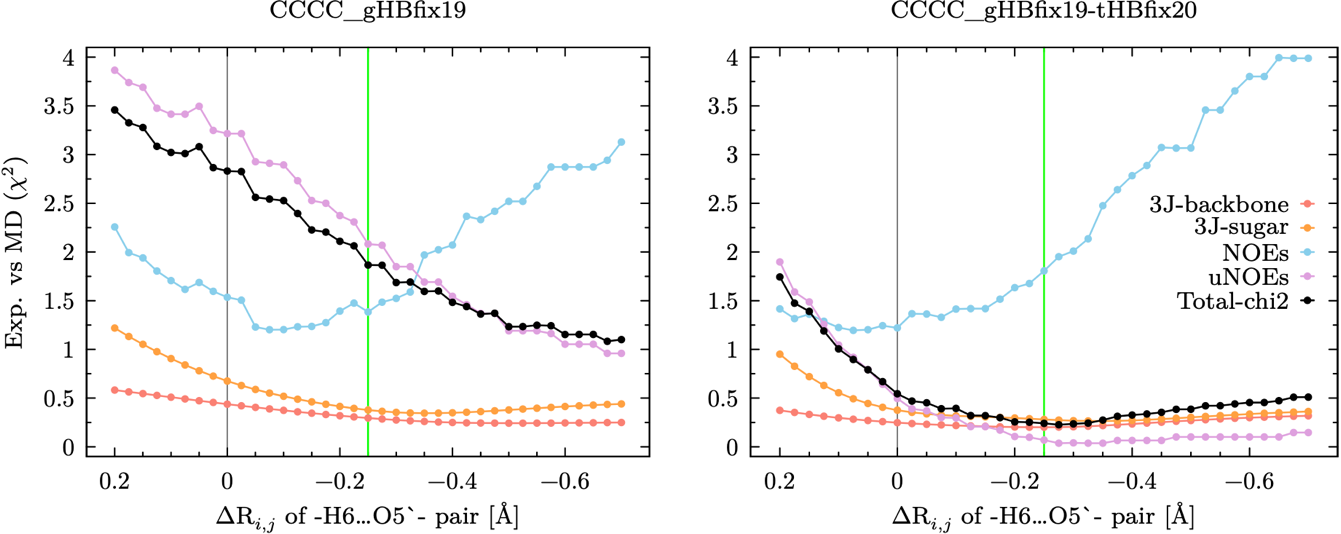

**Figure S2:** Effects of different settings of the NBfix_0BPh_ modification on agreement between simulations and NMR data as probed by reweighting of two r(CCCC) TN trajectories from REST2 simulations. Plots show dependence of NMR observables on changes of the Lennard-Jones *R_i,j_* parameter for pyrimidines, i.e., the distance between –H6…O5’– pairs, from the original (standard AMBER *ff*) values. See legend of Figure S1 for more details.

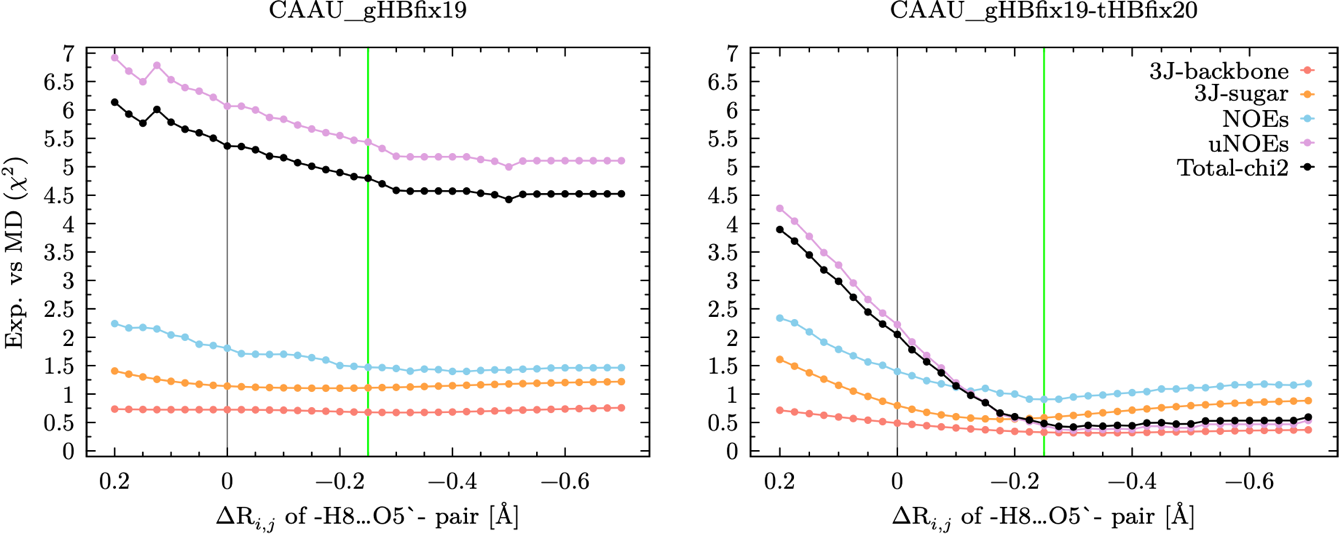

**Figure S3:** Effects of different settings of the NBfix_0BPh_ modification on agreement between simulations and NMR data as probed by reweighting of two r(CAAU) TN trajectories from REST2 simulations. Plots show dependence of NMR observables on changes of the Lennard-Jones *R_i,j_* parameter for purines, i.e., the distance between –H8…O5’– pairs, from the original (standard AMBER *ff*) values. See legend of Figure S1 for more details.

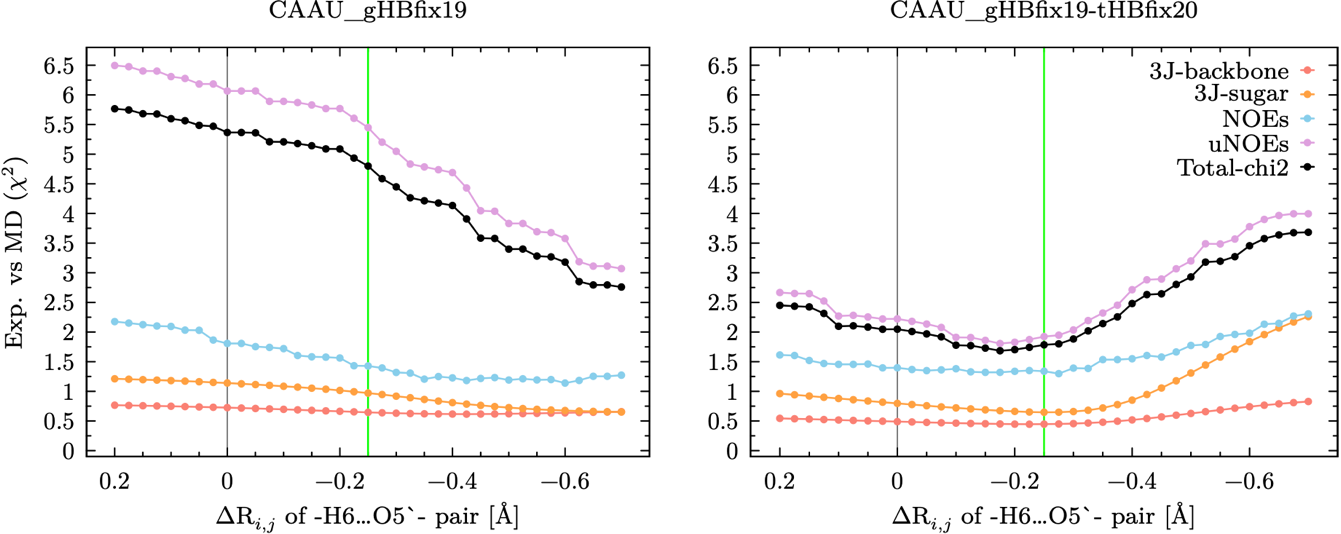

**Figure S4:** Effects of different settings of the NBfix_0BPh_ modification on agreement between simulations and NMR data as probed by reweighting of two r(CAAU) TN trajectories from REST2 simulations. Plots show dependence of NMR observables on changes of the Lennard-Jones *R_i,j_* parameter for pyrimidines, i.e., the distance between –H6…O5’– pairs, from the original (standard AMBER *ff*) values. See legend of Figure S1 for more details.

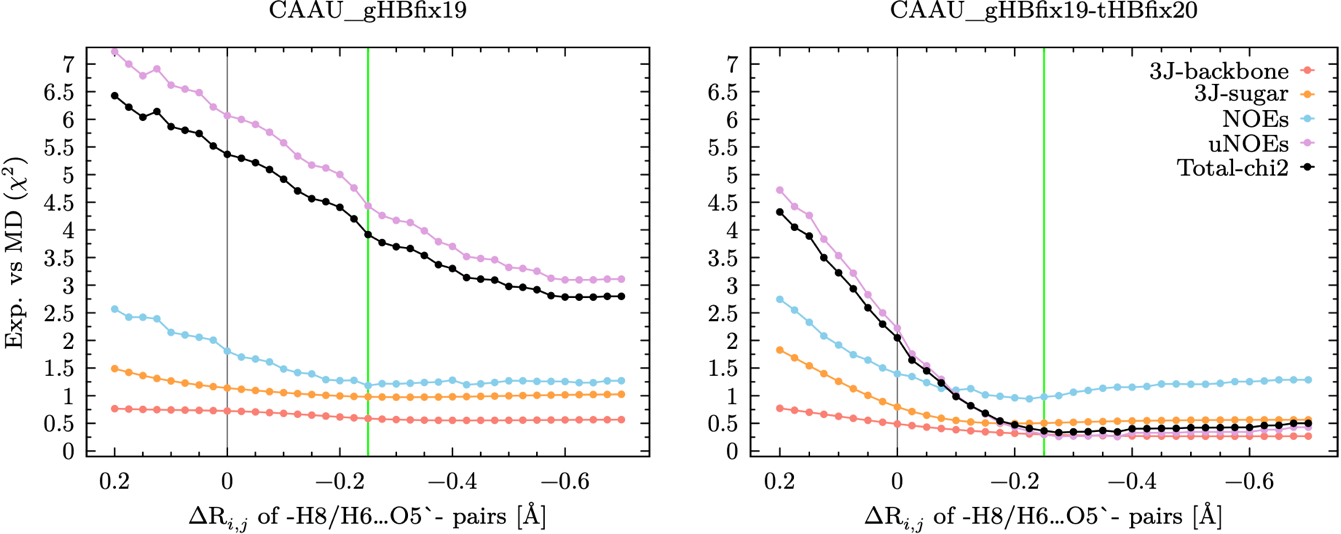

**Figure S5:** Effects of different settings of the NBfix_0BPh_ modification on agreement between simulations and NMR data as probed by reweighting of two r(CAAU) TN trajectories from REST2 simulations. Plots show dependence of NMR observables on changes of the Lennard-Jones *R_i,j_* parameter for both purines and pyrimidines, i.e., distances between –H8…O5’– pairs and –H6…O5’– pairs, respectively, from the original (standard AMBER *ff*) values. See legend of Figure S1 for more details.

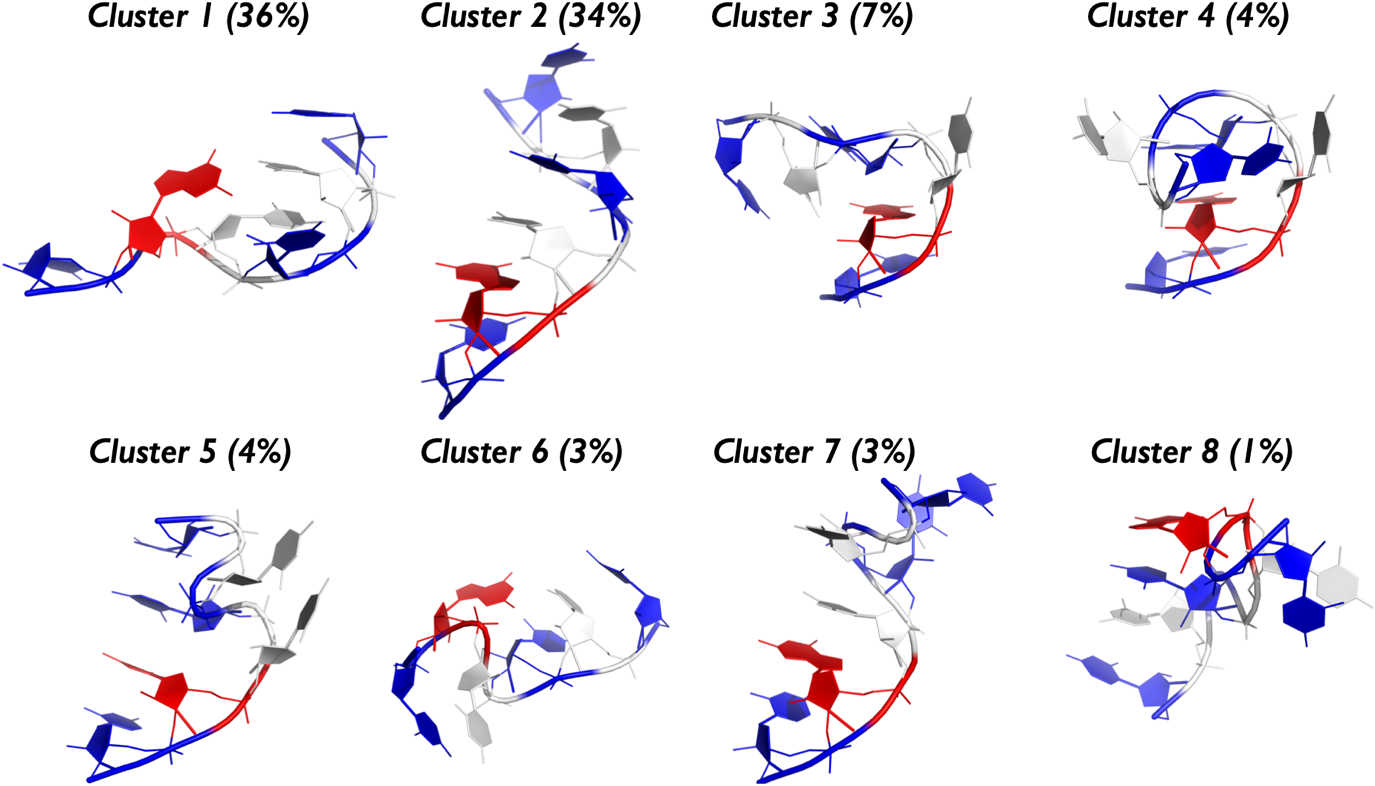

**Figure S6:** Tertiary structures of all populated clusters from r(UCUCGU) REST2 simulation with the OL3_CP_ RNA *ff*, gHBfix19 and tHBfix20 potentials and the NBfix_0BPh_ modification. C, G and U nucleotides are colored in white, red, and blue, respectively. H-atoms, ions and water molecules are not shown for clarity. Most populated conformers are loop-like state with G5C2 base pair and canonical A-form state. Population of unassigned structures is ~8%.

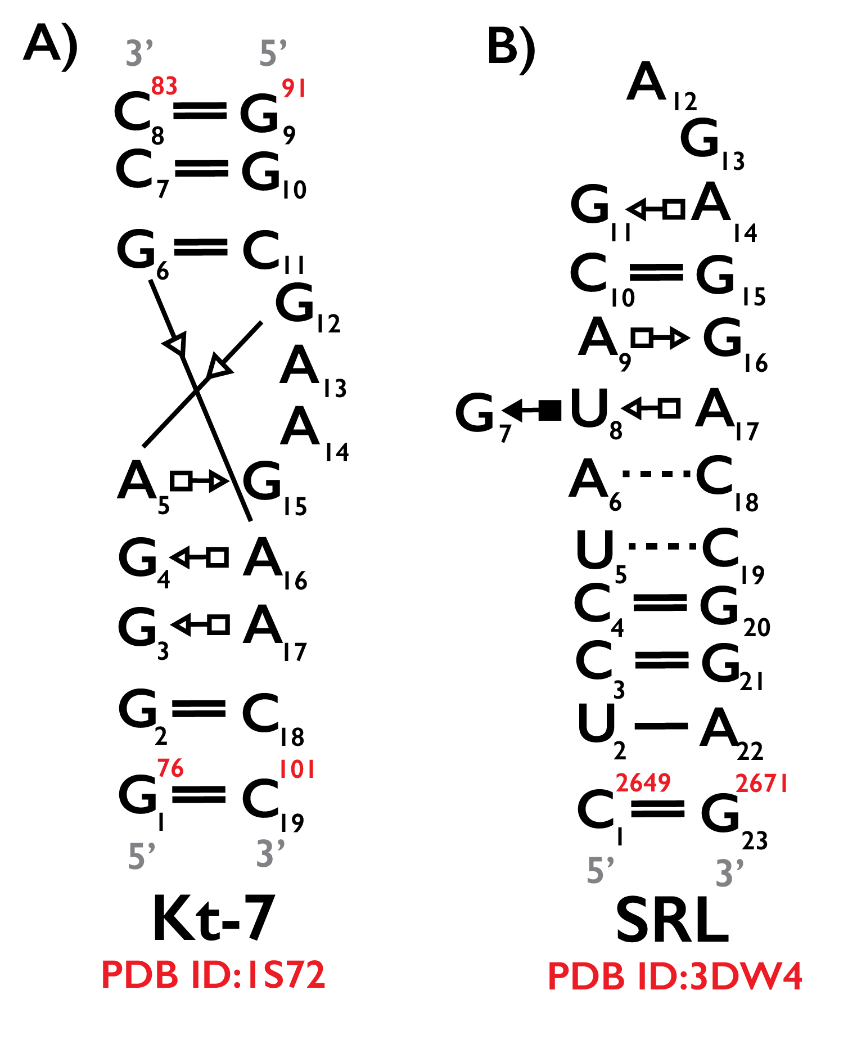

**Figure S7:** Secondary structure annotation following the Leontis−Westhof nomenclature^14^ for RNA kink-turn 7 (Kt-7, panel A) and Sarcin-Ricin loop (SRL, panel B) motifs. As starting coordinates for both those motifs were excised from larger experimental structures (see Methods in main text), we are showing both original (red) and relabeled (black) residue indexes. Starting coordinates are also attached (zipped PDB files).

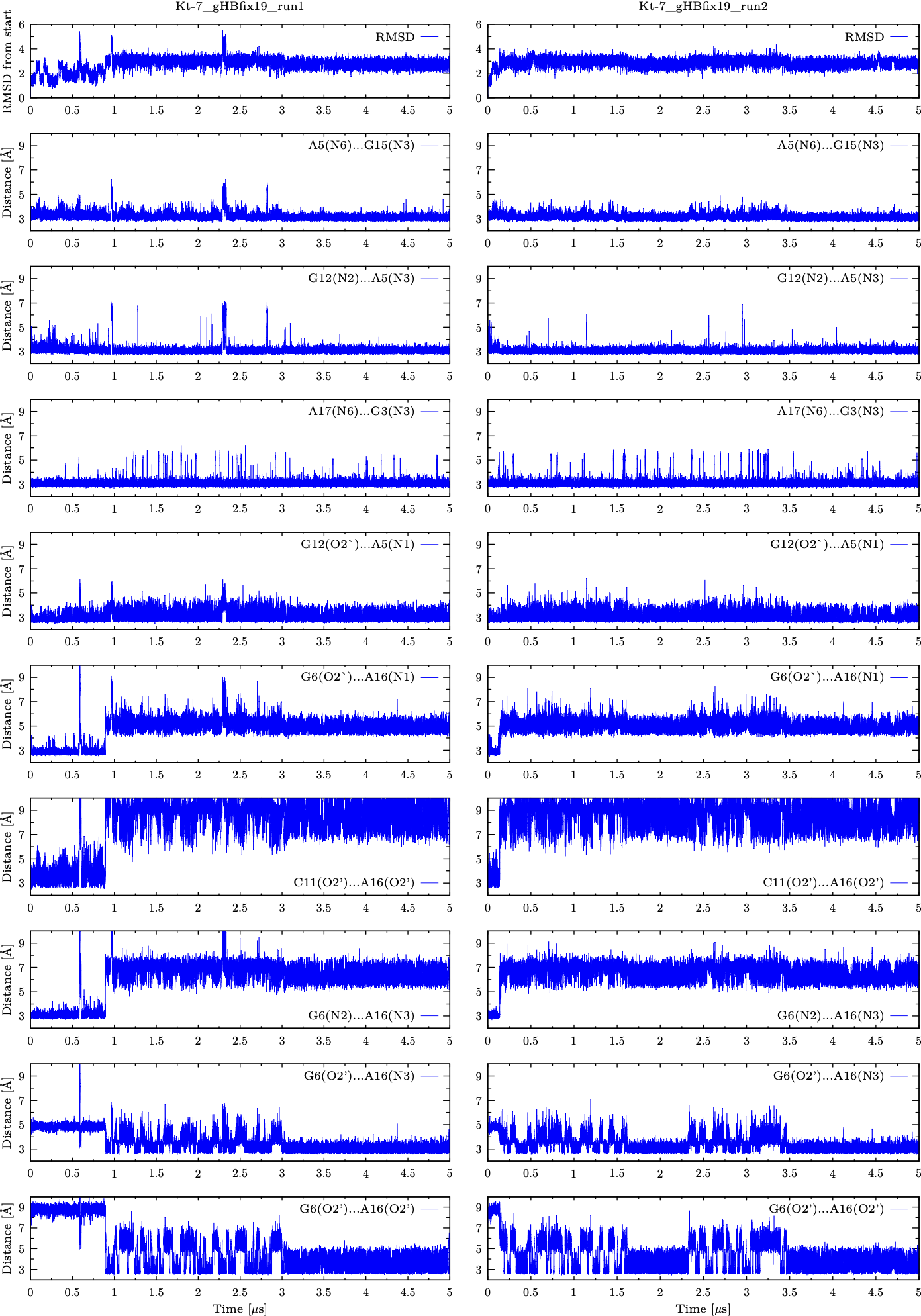

**Figure S8:** Structural analysis of two independent control standard MD simulations of the Kt-7 motif using OL3_CP_ *ff* with gHBfix19 potential (see Methods in main text and Table S1). Plots on the top are showing fluctuations of RMSD calculated over heavy atoms of all nucleotides during 5 μs-long unbiased MD simulations. Remaining plots are displaying fluctuations of key distances.

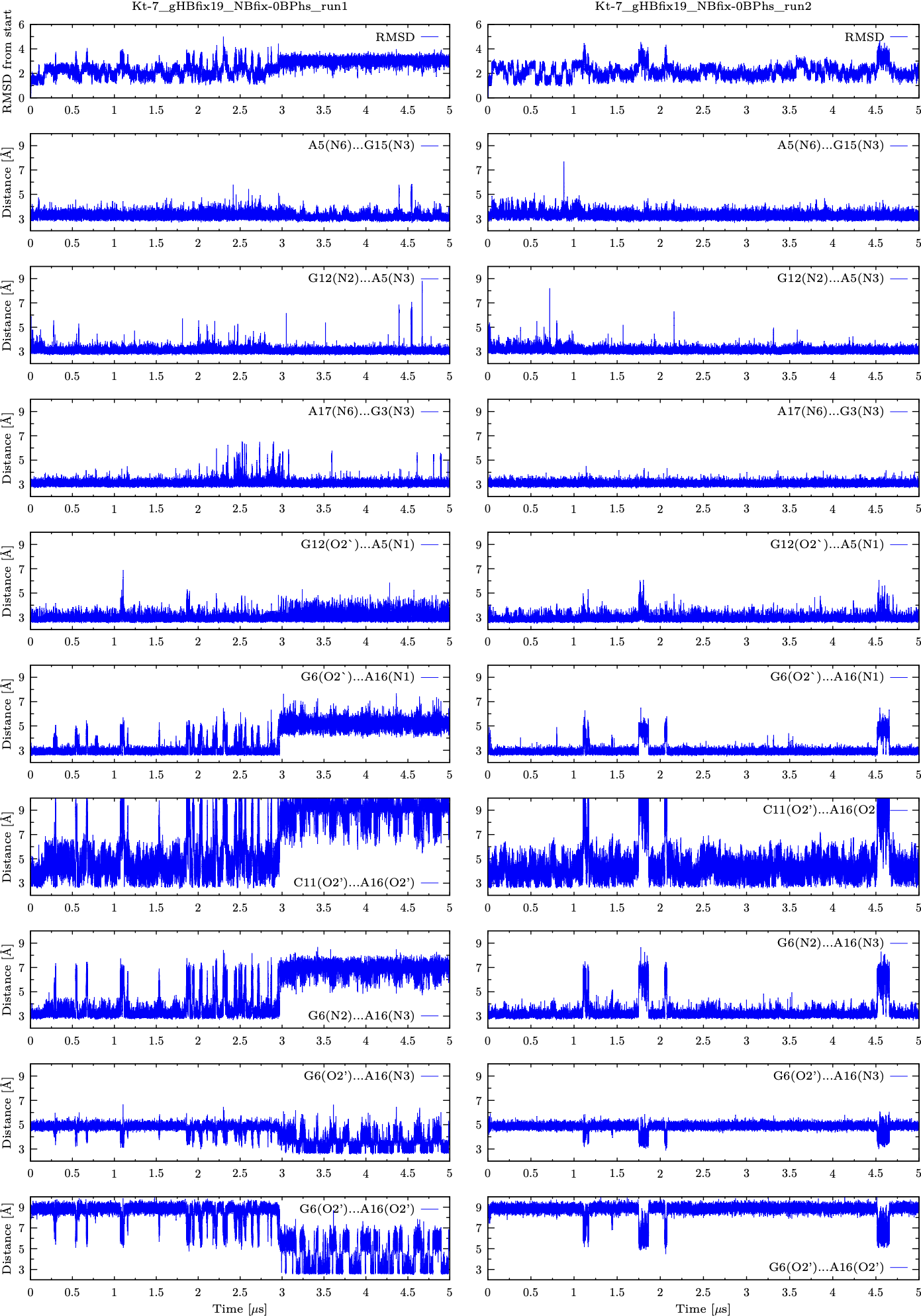

**Figure S9:** Structural analysis of two independent standard MD simulations of Kt-7 motif using OL3_CP_ *ff* with gHBfix19 potential and NBfix_0BPh_ modification (see legend of Figure S8 for more details).

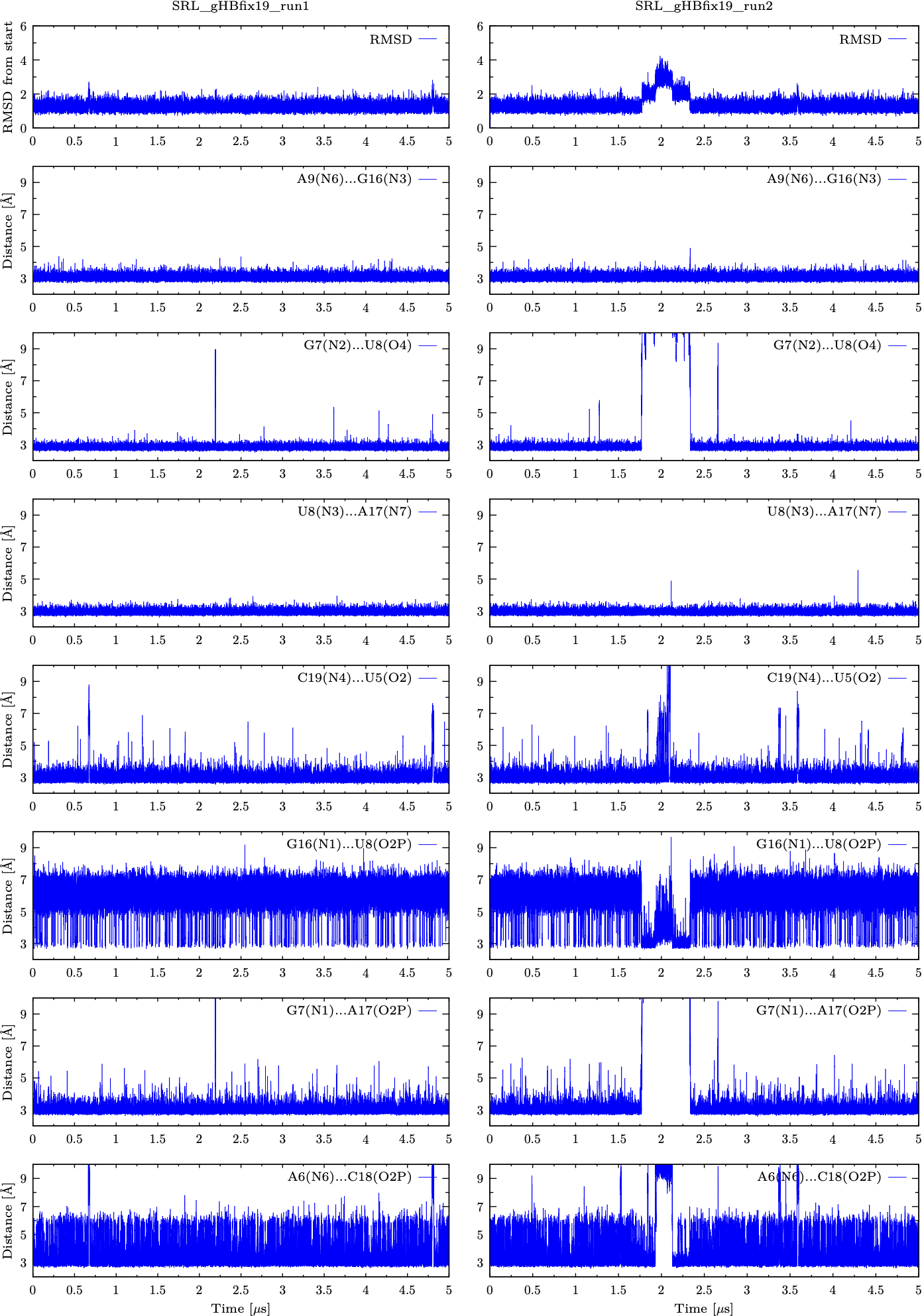

**Figure S10:** Structural analysis of two independent standard MD simulations of SRL motif using OL3_CP_ *ff* with gHBfix19 potential (see Methods in main text and Table S1). Plots on the top are showing fluctuations of RMSD calculated over heavy atoms of all nucleotides during 5 μs-long unbiased MD simulations. Remaining plots are displaying fluctuations of key distances.

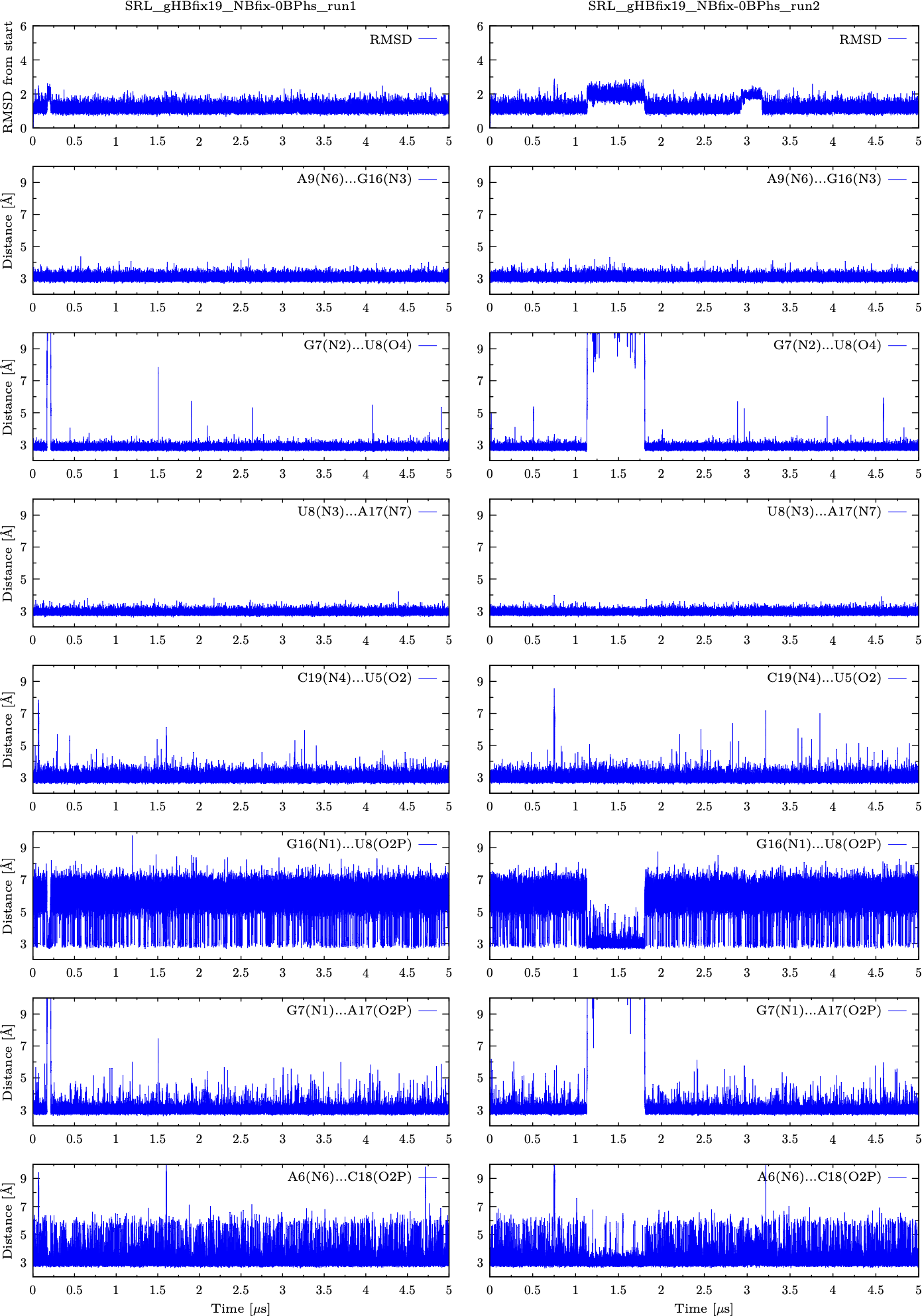

**Figure S11:** Structural analysis of two independent standard MD simulations of SRL motif using OL3_CP_ *ff* with gHBfix19 potential and NBfix_0BPh_ modification (see legend of Figure S10 for more details).

| 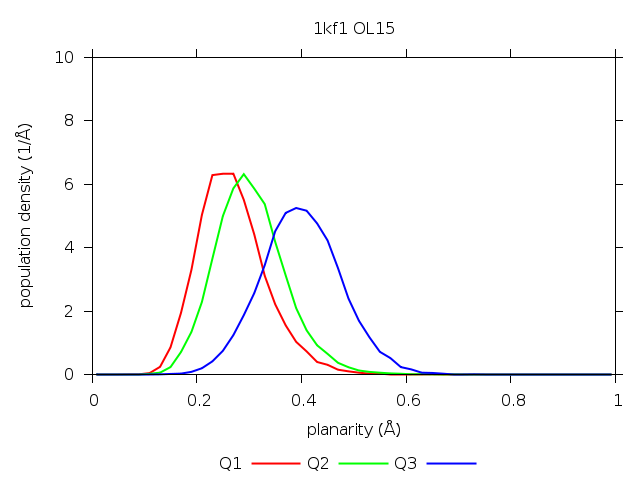 | 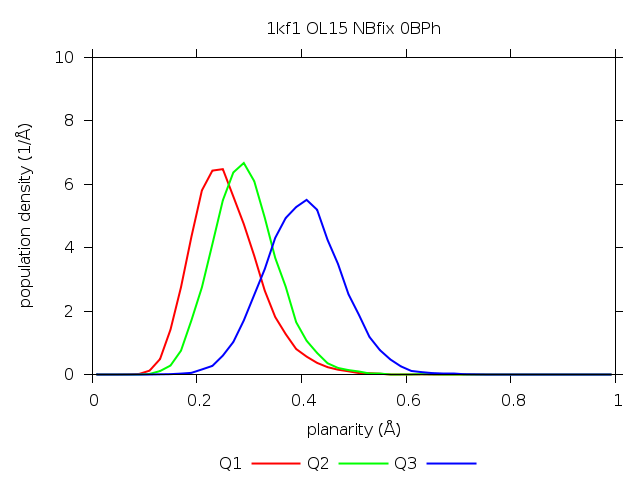 |
| --- | --- |
| 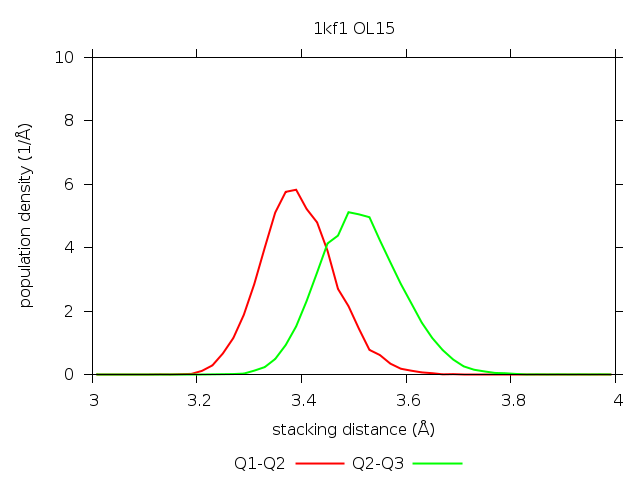 | 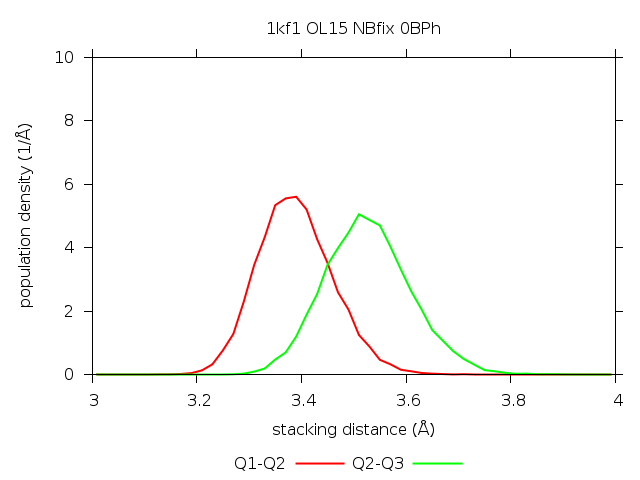 |
| 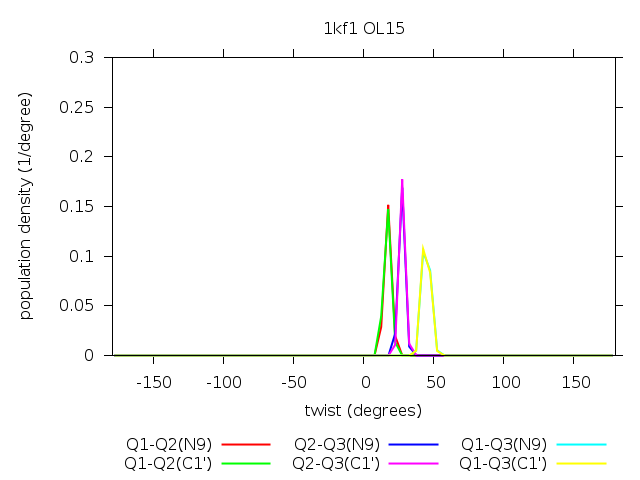 | 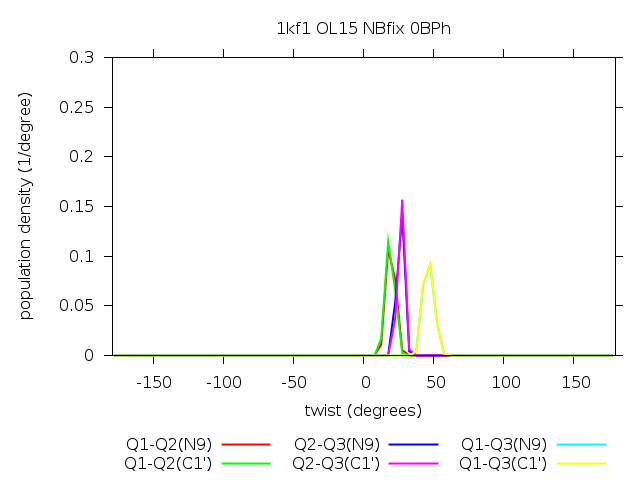 |

**Figure S12:** GQ structural parameters observed in the simulations of parallel-stranded 1KF1 GQ. The graphs show essentially identical (within the limits of sampling) distributions of quartet planarities (top row; Q1 – 5’-quartet, Q2 – middle quartet, Q3 – 3’quartet), stacking distances (middle row) and helical twists (bottom row; measured either by the position of C1’ atoms or N9 atoms) between adjacent quartets in the standard OL15 *ff* (control; left column) and the version with added NBfix_0BPh_ term (right column). Planarity of a quartet was defined as the root mean square deviation of the distances of non-hydrogen atoms to the best-fit quartet plane. Stacking distance between two quartets was calculated as a sum of distances of the two quartet centers to the quartets' best-fit planes average.^15^

| 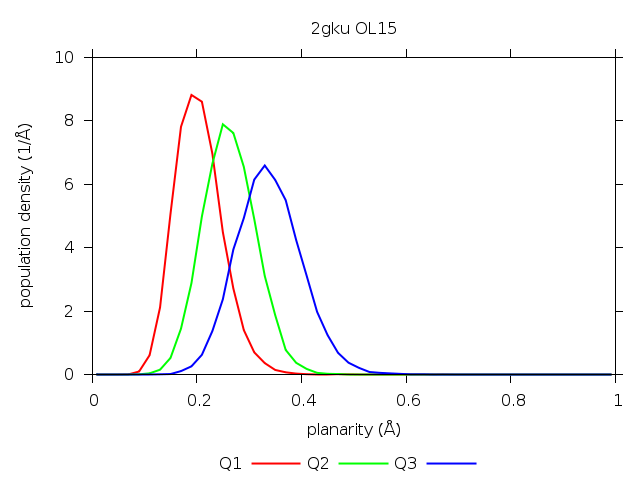 | 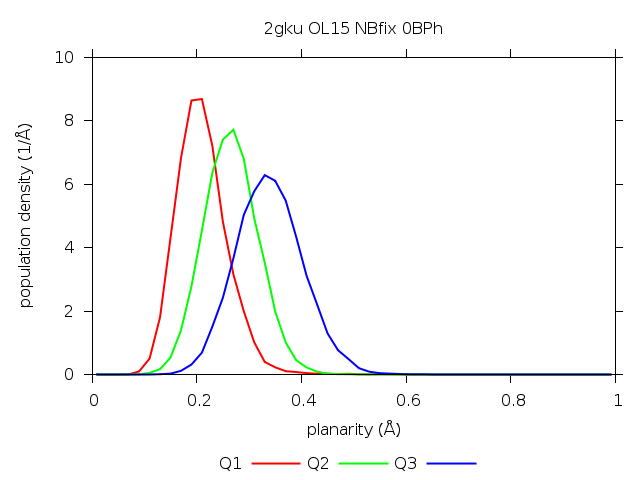 |
| --- | --- |
| 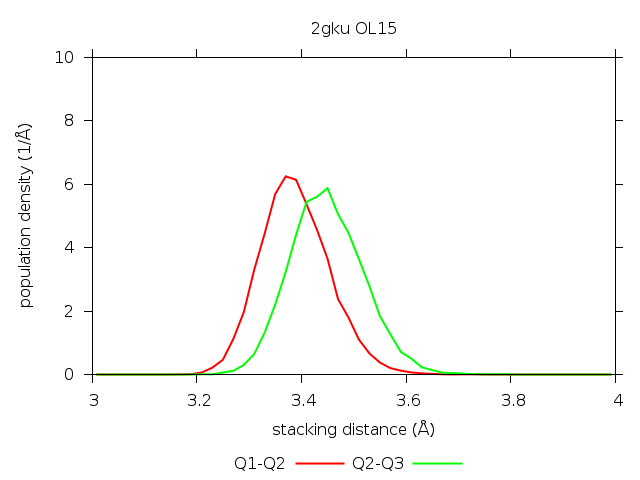 | 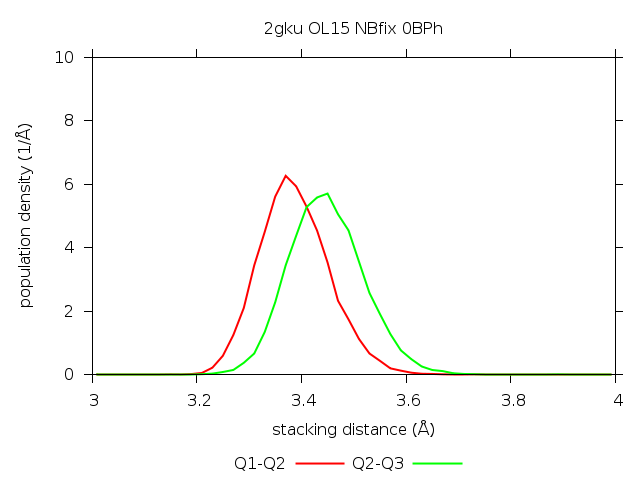 |
| 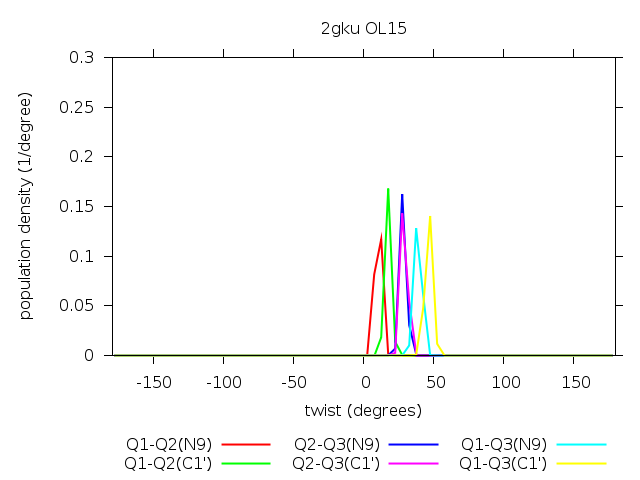 | 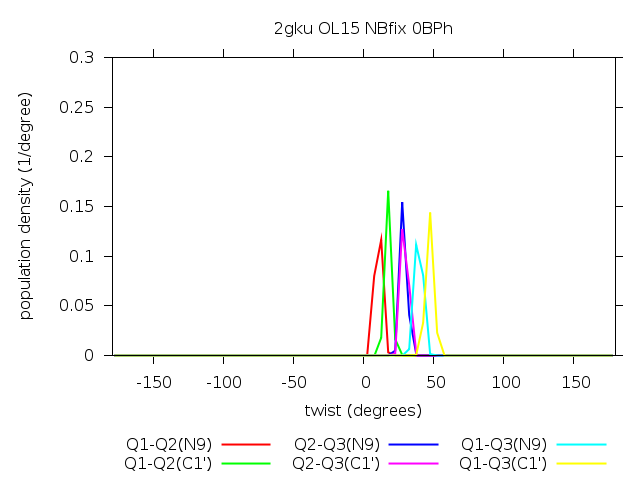 |

**Figure S13:** GQ structural parameters observed in the simulations of (3+1) hybrid 2GKU GQ. The graphs show identical (within the limits of sampling) outcome between the standard OL15 *ff* (control; left column) and the version with NBfix_0BPh_ term (right column). See legend of Figure S12 for more details.

Research Support, U.S. Gov't, Non-P.H.S.

Research Support, U.S. Gov't, P.H.S. DOI: 10.1017/s1355838201002515.

(15) Zhang, Z., Stadlbauer, P., Mlýnský, V., Krepl, M., and Šponer J. Tetrameric Parallel Guanine-Quadruplex Folding Pathways Unveiled by Steered Molecular Dynamics Simulations. *in preparation* **2023**.
